# Accurate promoter and enhancer identification in 127 ENCODE and Roadmap Epigenomics cell types and tissues by GenoSTAN

**DOI:** 10.1101/041020

**Authors:** Benedikt Zacher, Margaux Michel, Björn Schwalb, Patrick Cramer, Achim Tresch, Julien Gagneur

## Abstract

Accurate maps of promoters and enhancers are required for understanding transcriptional regulation. Promoters and enhancers are usually mapped by integration of chromatin assays charting histone modifications, DNA accessibility, and transcription factor binding. However, current algorithms are limited by unrealistic data distribution assumptions. Here we propose GenoSTAN (Genomic STate ANnotation), a hidden Markov model overcoming these limitations. We map promoters and enhancers for 127 cell types and tissues from the ENCODE and Roadmap Epigenomics projects, today’s largest compendium of chromatin assays. Extensive benchmarks demonstrate that GenoSTAN consistently identifies promoters and enhancers with significantly higher accuracy than previous methods. Moreover, GenoSTAN-derived promoters and enhancers showed significantly higher enrichment of complex trait-associated genetic variants than current annotations. Altogether, GenoSTAN provides an easy-to-use tool to define promoters and enhancers in any system, and our annotation of human transcriptional cis-regulatory elements constitutes a rich resource for future research in biology and medicine.

## Introduction

Transcription is tightly regulated by cis-regulatory DNA elements known as promoters and enhancers. These elements control development, cell fate and may lead to disease if impaired. A promoter is functionally defined as a region that regulates transcription of a gene, located upstream and in close proximity to the transcription start sites (TSSs) 1. In contrast, an enhancer was originally functionally defined as a DNA element that can increase expression of a gene over a long distance in an orientation-independent fashion relative to the gene 2. The functional definition of enhancers and promoters leads to practical difficulties for their genome-wide identification because the direct measurement of the regulatory activity of genomic regions is hard, with current approaches leading to contradicting results [3, 4, 5].

Since the direct measurement of cis-regulatory activity is challenging, a biochemical characterization of the chromatin at these elements based on histone modifications, DNA accessibility, and transcription factor binding has been proposed [6, 7, 8, 9, 10]. This approach leverages extensive genome-wide datasets of chromatin-immunoprecipitation followed by sequencing (ChIP-Seq) of transcription factors (TFs), histone modifications, or Cap analysis gene expression (CAGE) that have been generated by collaborative projects such as ENCODE [11, 12], NIH Roadmap Epigenomics 13, BLUEPRINT 14 and FANTOM [15, 16].

In this context, the computational approaches employed to classify genomic regions as enhancers or promoters play a decisive role [6, 10]. As the experimental data are heterogeneous, we generally refer to them as tracks. Several studies used supervised learning techniques to predict enhancers based on tracks such as histone modifications or P300 binding (e.g. [17, 18, 19, 20]). However, a training set of validated enhancers is needed in this case, which is hard to define since only few enhancers have been validated experimentally so far and these might be biased towards specific enhancer subclasses. Alternatively, unsupervised learning algorithms were developed to identify promoters and enhancers from combinations of histone marks and protein-DNA interactions alone [8, 21, 22, 9, 23, 24, 13, 11]. These unsupervised methods perform genome segmentation, i.e. they model the genome as a succession of segments in different chromatin states defined by characteristic combinations of histone marks and protein-DNA interactions found recurrently throughout the genome. All popular genome segmentations are based on hidden Markov models 25. However, these methods differ in the way the distribution of ChIP-seq signals for each chromatin state is modeled. ChromHMM [8, 26, 21], one of the two methods applied by the ENCODE consortium, requires binarized ChIP-seq signals that are then modeled with independent Bernoulli distributions. Consequently, the performance of ChromHMM highly depends on the non-trivial choice of a proper binarization cutoff. Moreover, quantitative information is lost with this approach, which is especially important for distinguishing promoters from enhancers since these elements are both marked with H3K4me1 and H3K4me3, but at different ratios 27. Segway [9, 22], the other method applied by the ENCODE consortium, uses independent Gaussian distributions of log-transformed and smoothed ChIP-seq signal. Although Segway preserves some quantitative information, the transformation of the original count data leads to variance estimation difficulties for very low counts. Therefore, Segway further makes the strong assumption that all tracks have the same variance. Recently, EpicSeg 28 used a negative multinomial distribution to directly model the read counts without the need for data transformations. However, similar to the variance model of Segway, the EpicSeg model leads to a common dispersion (the parameter adjusting the variance of the negative multinomial) for all tracks. Moreover, EpicSeg does not provide a way to correct for sequencing depth, which makes the application to data sets with multiple cell types with varying library sizes difficult. Also, EpicSeg has been applied only to three cell types so far 28. These methods not only differ in their modeling assumptions but also lead to very different results. In the K562 cell line for instance, ChromHMM identified 22,323 enhancers 11, Segway 38,922 enhancers 11, and EpicSeg 53,982 enhancers 28. Altogether, improved methods and detailed benchmarkings are required for a reliable annotation of transcriptional cis-regulatory elements.

Here we propose a new unsupervised genome segmentation algorithm, GenoSTAN (***Geno***mic ***St***ate ***An***notation from sequencing experiments), which overcomes limitations of current state-of-the-art models. GenoSTAN learns chromatin states directly from sequencing data without the need of data transformation, while still having track-specific variance models. We applied GenoSTAN to a total of 127 cell types and tissues covering 16 datasets of ENCODE and all 111 datasets of the Roadmap Epigenomics project as well as another ENCODE ChIP-seq dataset for the K562 cell line. GenoSTAN consistently performed better when benchmarked against Segway, ChromHMM and EpicSeg segmentations using independent evidence for activity of promoter and enhancer regions. Co-binding analysis of TFs reveals that promoters and enhancers both shared the Polymerase II core transcription machinery and general TFs, but they are bound by distinct TF regulatory modules and differ in many biophysical properties. Moreover, GenoSTAN enhancer and promoter annotations had a higher enrichment for complex trait-associated genetic variants than previous annotations, demonstrating the advantage of GenoSTAN and our chromatin state map to understand genotype-phenotype relationships and genetic disease.

## Results

### Modeling of sequencing data with Poisson-lognormal and negative binomial distributions

We developed a new genomic segmentation algorithm, GenoSTAN, which implements hidden Markov models with more flexible multivariate count distributions than previously proposed. Specifically, GenoSTAN supports two multivariate discrete emission functions, the Poisson-lognormal distribution and the negative binomial distribution. For the sake of reducing running time, the components of these multivariate distributions are assumed to be independent. However, the variance is modeled separately for each state and each track, which provides a more realistic variance model than current approaches. To be applicable to data sets with replicate experiments or multiple cell types, GenoSTAN corrects for different library sizes of experiments (Methods). All parameters are learnt directly from the data, leaving the number of chromatin states as the only parameter to be manually set. We provide an efficient implementation of the Baum-Welch algorithm for inference of model parameters, which can be run in a parallelized fashion using multiple cores. The method is implemented as part of our prevsiouy published R/Bioconductor package STAN 29, which is freely available from bioconductor.org/. Altogether, GenoSTAN uniquely combines flexible count distributions, short running times, and minimal number of manually entered parameters (Figure 1A).

**Figure 1:**
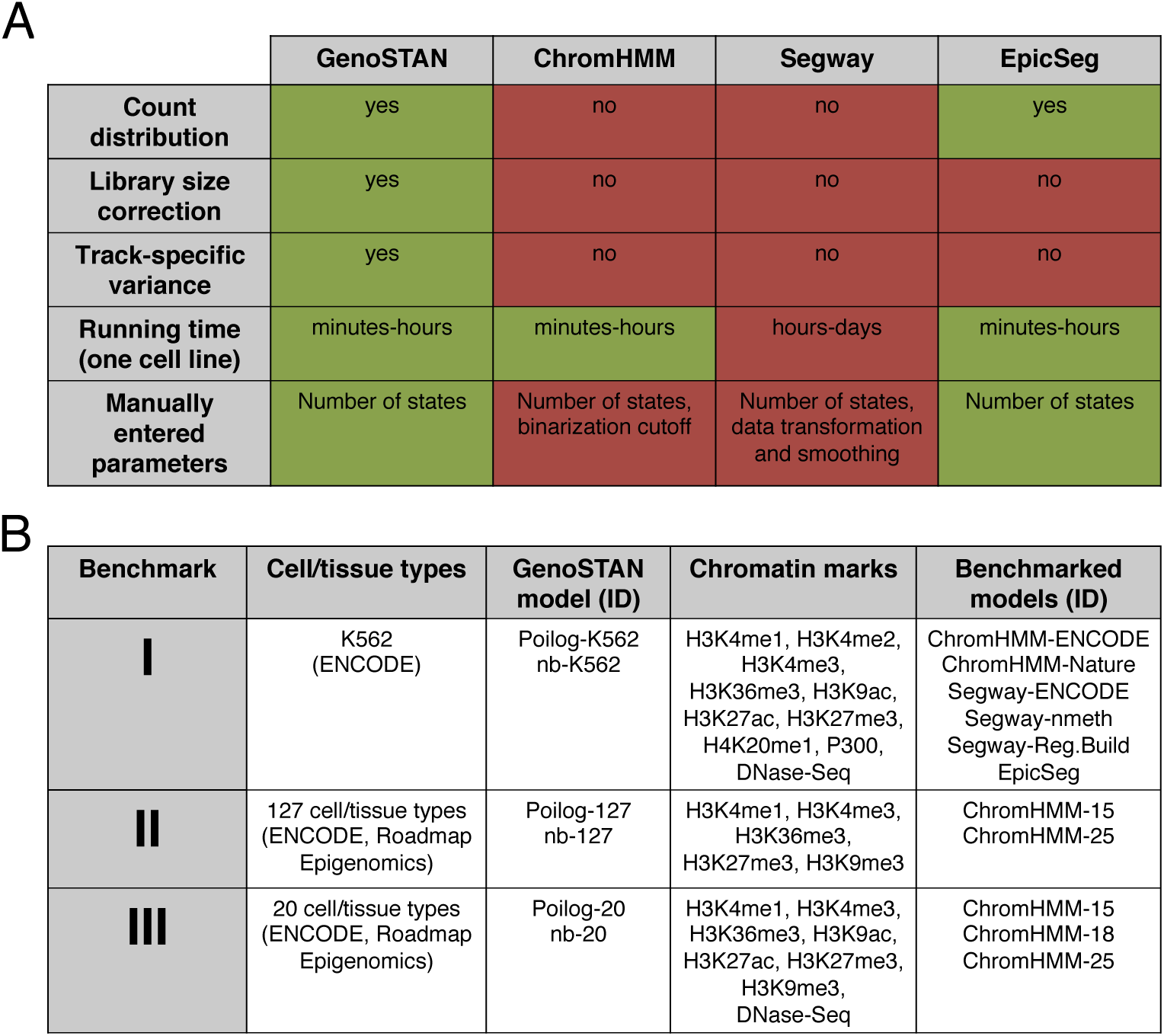
Overview of chromatin state annotation methods and study design. (A) Comparison of features of GenoSTAN against three previous chromatin state annotation algorithms. (B) Description of the three benchmark sets used in this study. GenoSTAN is benchmarked against published chromatin state annotations using ChromHMM (’ChromHMM-ENCODE’ [11, 22], ‘ChromHMM-Nature’ 21, ‘ChromHMM-15’, ‘-18’ and ‘-25’ 13), Segway (’Segway-ENCODE’ [11, 22], ‘Segway-nmeth’ 9 and ‘Segway-Reg.Build’ 23) and EpicSeg 28.

We performed an extensive benchmarking of GenoSTAN against alternative methods (Figure 1B). Benchmark I compares GenoSTAN and the three alternative methods for the K562 cell line. K562 is a major model system to study human transcription and the ENCODE cell line with the largest number of experiments 11. Moreover, all three other methods (ChromHMM, Segway, and EpicSeg) had been run by their own authors for K562, so that the algorithm parameters for these annotations can be assumed to have been set at best expert knowledge. Benchmark II compares methods for all 127 cell types and tissues provided by ENCODE and Roadmap Epigenomics. This is the largest benchmark dataset of all three we considered but also the one with the least number of common tracks (5 chromatin marks: H3K4me1, H3K4me3, H3K36me3, H3K27me3, and H3K9me3, Figure 1B). Benchmark III compares results for a subset of 20 ENCODE and Roadmap Epigenomics cell types and tissues which had moreover H3K27ac, H3K9ac ChIP-seq and DNAse I hypersensitivity data (DNAse-Seq). This distinction allowed to provide more accurate annotations for the better characterized cell types and tissues. For benchmark II and benchmark III only annotations for the method ChomHMM were available to compare against GenoSTAN annotation. Figure 1B lists all model names and studies that were considered.

### Benchmark I: Improved chromatin state annotation in human K562 cells

We first fitted two GenoSTAN models, one with Poisson-lognormal emissions (henceforth referred to as GenoSTAN-Poilog-K562 model) and one with negative binomial emissions (GenoSTAN-nb-K562 model) to a dataset of ChIP-seq data of 9 histone modifications, of the histone acetyltransferase P300, and DNA accessibility (by DNase-Seq) data for the K562 cell line at 200 bp binning resolution (Methods). As pointed out by others [8, 9], there is no purely statistical criterion for choosing the number of states from the data of practical usage in such a setting. In practice, the number of states is manually defined by trading off goodness of fit against interpretability of the model [8, 9, 29]. For GenoSTAN-Poilog-K562, we used 18 chromatin states. For GenoSTAN-nb-K562, we used 23 states, since lower state numbers did not provide enough resolution to give a fine-grained map of chromatin states on this data set (Methods). Figure 2A compares the two GenoSTAN segmentations to segmentations from other studies using ChromHMM, Segway and EpicSeg on a region containing the TAL1 gene together with three known enhancers [28, 22, 11, 30].

**Figure 2:**
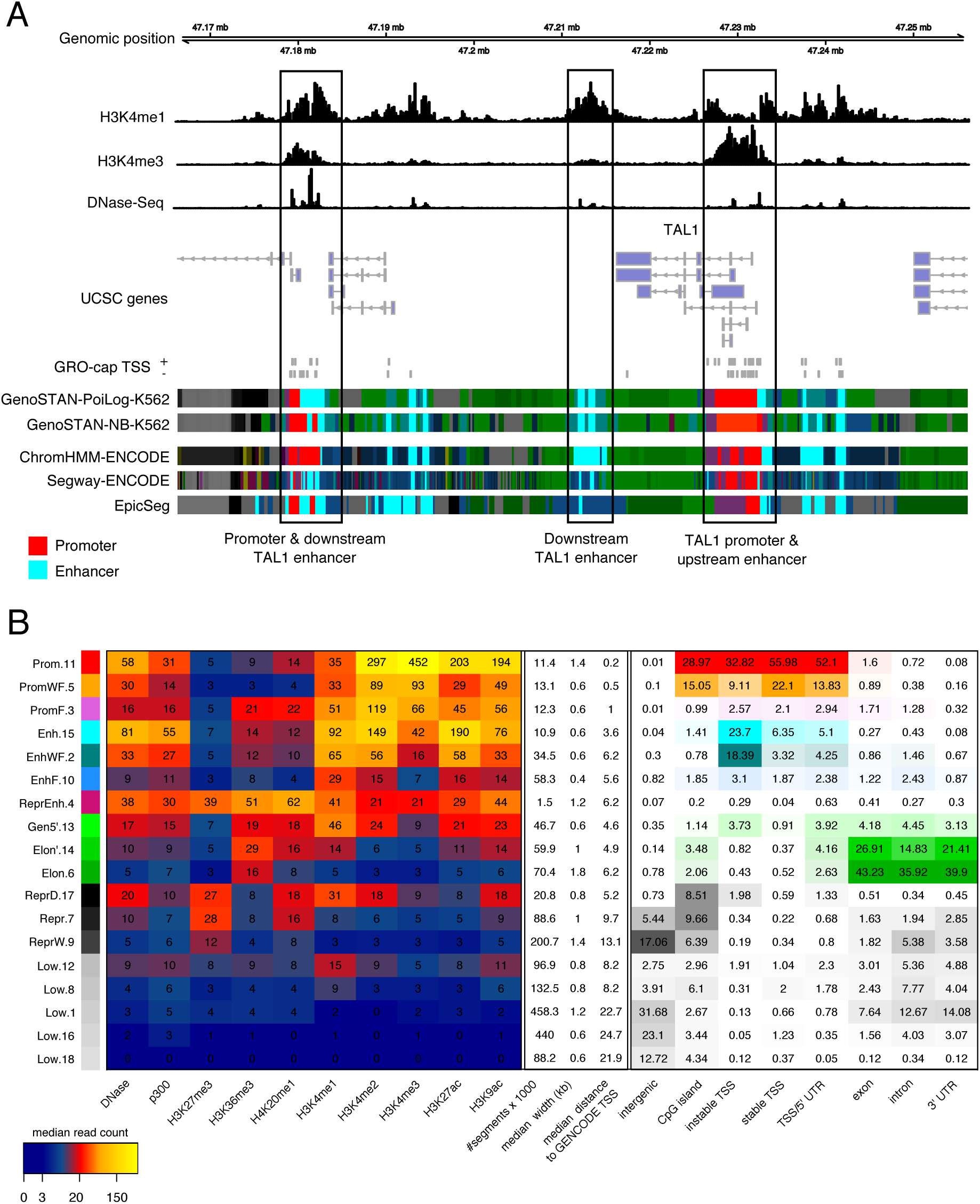
Chromatin states fitted on Benchmark I using GenoSTAN. (A) GenoSTAN segmentations are shown with published segmentations using ChromHMM-ENCODE 11, Segway-ENCODE 11 and EpicSeg 28 at the TAL1 gene and three known enhancers. GenoSTAN-Poilog-K562 correctly recalls all known promoter and enhancer regions. GenoSTAN-nb-K562 misses the upstream enhancer. ChromHMM-ENCODE misclassifies most of the downstream enhancer region as promoter. (B) Median read coverage of GenoSTAN-Poilog-K562 chromatin states (left), their number of annotated segments in the genome, their median width and distance to the closest GENCODE TSS (middle). The right panel shows recall of genomic regions by chromatin states.

#### Chromatin states recover biologically meaningful features

In order to assign biologically meaningful labels to each state of the Poilog-K562 and of the nb-K562 GenoSTAN models, we investigated their read coverage distributions and overlapped the occurrence of a state in the genome with known genomic features. In line with previous studies, this led to the definition of promoter, enhancer, repressed, actively transcribed and low coverage states [28, 21, 22]. The median read coverage in state segments and genomic distributions were very similar for both the Poilog-K562 and the nb-K562 models (Figure 2B, Supplementary Figure 1). Promoter states were characterized by a low (< 1) H3K4me1/H3K4me3 ratio, in contrast to enhancer states which showed a high ratio (> 1). Further, P300 levels were roughly two-fold higher in the enhancer state, which is in accordance with previous observations [27, 31, 32]. Promoter (Prom) states were located close to annotated GENCODE TSSs 33, with a median distance of 220 bp for GenoSTAN-Poilog-K562 and 400 bp for GenoSTAN-nb-K562 model. Enhancer states (Enh) on the other hand were located further away from TSSs, with a median distance of 3.6 kb (Poilog-K562) (respectively 5.8 kb for nb-K562, Figure 2B, Supplementary Figure 1). Promoter and enhancer states also differed in their DNA sequence features. 45% of CpG islands were located within promoter states (strong, weak and promoter flanking states) in both models, but only 3% in enhancer states (strong, weak, enhancer flanking states, Figure 2B, Supplementary Figure 1). While promoter states mostly recovered stable TSSs, enhancer states were located at unstable TSSs (GRO-cap TSSs that are not recovered by the GENCODE annotation), which supports previous findings 34. Furthermore, both models (GenoSTAN-nb-K562 and GenoSTAN-Poilog-K562) contained 3 states, that we classified as “actively transcribed states”, which were characterized by high values of H3K36me3 and overlap with UTRs, introns and exons. Two out of three “transcribed” states were also enriched in promoter associated marks (H3K4me1-3, H3K27ac, H3K9ac) and H4K20me1 and thus represented 5’ transitions in transcription. Moreover, both models fitted four repressed states showing high read coverage of H3K27me3. Two of these states also exhibited high DNase-Seq and promoter/enhancer associated histone modification signals, suggesting that these states might reflect repressed regulatory regions (ReprEnh, ReprD). These elements were distal to annotated GENCODE TSSs (median distance: 5.2-11.8 kb). ReprEnh states were also enriched in P300 and recovered 0.2% of CpG islands, while ReprD states had lower P300 levels and recovered 8-9% of CpG islands in the genome (Figure 2B, Supplementary Figure 1). The remaining states exhibited low coverage in chromatin marks and therefore were labeled as “low” states. Altogether, GenoSTAN accurately recovered many features of known chromatin states and provided a high resolution map of these in K562.

#### High variation of enhancer predictions between chromatin state annotations of different studies

To assess the consistency of promoter and enhancer predictions across studies, we compared the GenoSTAN segmentations to other published segmentations in K562 by ChromHMM (’ChromHMM-ENCODE’ [11, 22] and ‘ChromHMM-Nature’ 21), Segway (’Segway-ENCODE’ [11, 22], ‘Segway-nmeth’ 9 and ‘Segway-Reg.Build’ 23) and EpicSeg 28. We computed pairwise Jaccard indices (the ratio of the number of common elements over all elements predicted by two methods) of promoter and enhancer states to quantify the agreeement between the predictions of the different studies (Supplementary Figure 2). Promoter state annotations generally agreed well (median Jaccard-Index: 0.78). However, enhancer prediction varied more (median Jaccard Index: 0.48), suggesting that enhancers are more difficult to annotate. This variation of enhancer calls was also reflected in the different numbers of annotated enhancer segments, which had been shown to vary greatly between different prediction methods 17. The number of enhancer segments ranged from 10,932 segments in GenoSTAN-Poilog-K562 to 80,043 segments in one Segway annotation 9 (Supplementary Table 1). Therefore, a thorough assessment of these predictions was necessary to provide a robust and accurate prediction of these elements.

#### Comparison of GenoSTAN with published chromatin state annotations

In order to benchmark the different segmentations, we used independent data including evidence of transcriptional activity (GRO-cap TSSs 34), of transcription factor binding (ENCODE high occupancy target, or HOT regions 12, and ENCODE TF binding sites 11), and of cis-regulatory activity (enhancer activity assessed by reporter assays 4), which are all expected to be characteristics of promoters and enhancers. Transcription initiation activity is not only the hallmark of promoters, but also of enhancers [15, 16, 34, 35]. To benchmark the predictions using evidence for transcription, we used published data from a protocol called GRO-cap 34, a nuclear run-on protocol, which very sensitively maps transcription start sites genome-wide. To this end, we sorted for each method chromatin states by their overlap with GRO-cap TSSs by decreasing precision. Starting with the most precise state (i.e. highest overlap with TSSs) we calculated cumulative recall and false discovery rate (FDR) by subsequently adding states with decreasing precision (Figure 3A). GenoSTAN-Poilog-K562 had the highest recall and the lowest FDR (Methods, Figure 3A). GenoSTAN-nb-K562 performed similar to other segmentations (Segway-Reg. Build, ChromHMM-ENCODE). In particular, 94% of GenoSTAN-Poilog-K562 promoters (Prom.11) and 81% of its enhancer regions (Enh.15) overlapped with GRO-cap TSSs. This compares to 85% (Prom.16) and 65% (Enh.6) of GenoSTAN-nb-K562 and 89% (Tss) and 52% (Enh) of ChromHMM-ENCODE promoter and enhancer regions. Interestingly, the two ChromHMM segmentations (ChromHMM-ENCODE [22, 11], ChromHMM-Nature 21) had very different accuracies for TSSs, which might be due to different data sets or cutoffs used for ChIP-seq binarization in the two studies. In contrast, the overall accuracy of the Segway annotations was comparable across studies. This comparison shows that GenoSTAN chromatin state annotation identifies putative promoters and enhancers which show transcriptional activity more frequently than previous annotations.

**Figure 3:**
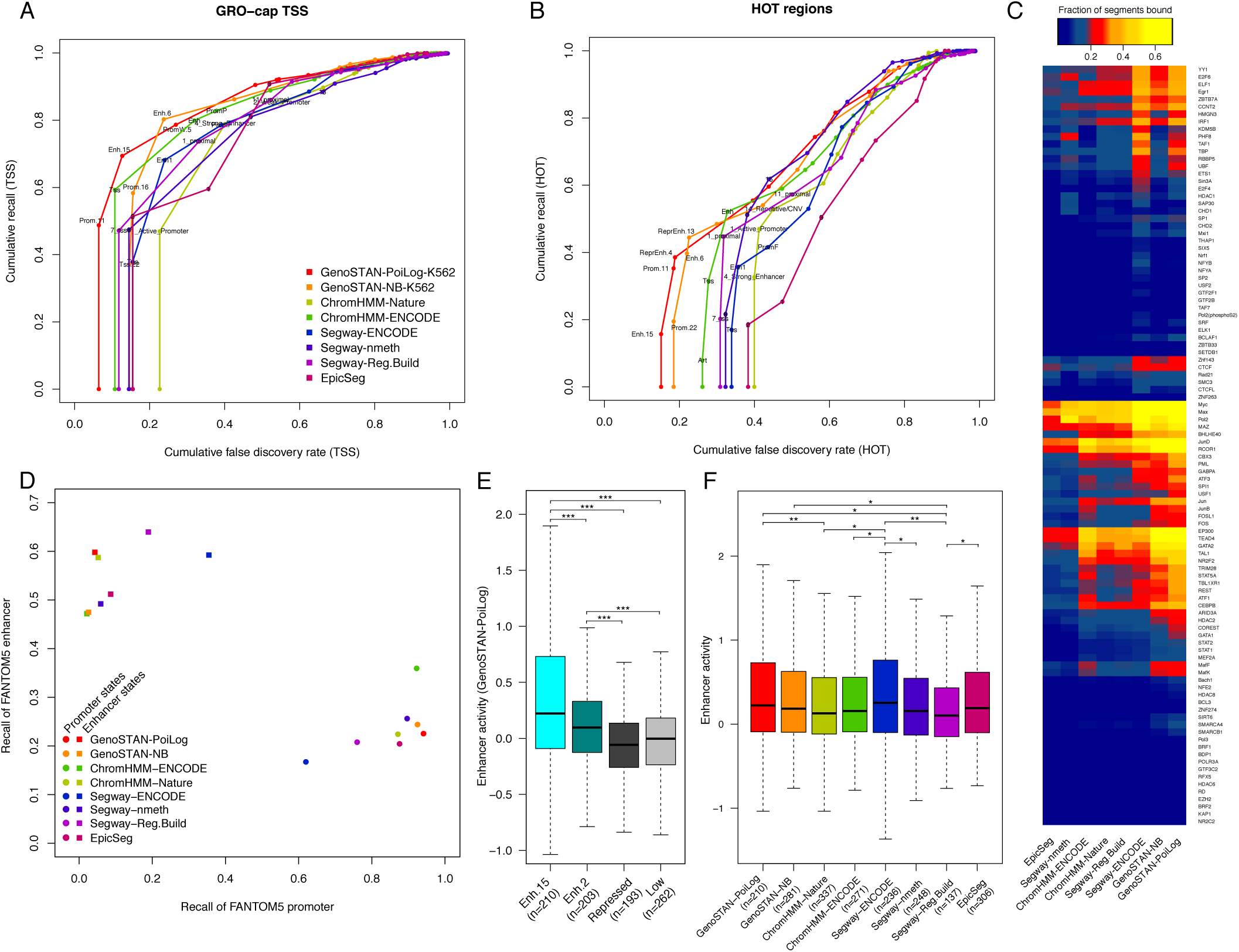
Comparison of GenoSTAN to other published segmentations on benchmark set I. (A) Performance of chromatin states in recovering GRO-cap transcription start sites. Cumulative FDR and recall are calculated by subsequently adding states (in order of increasing FDR). (B) The same as in (A) for ENCODE HOT regions. (C) The fraction of predicted enhancer segments bound by individual TFs is shown for different studies. GenoSTAN enhancers are more frequently bound by TFs than those from other studies. (D) Recall of FANTOM5 promoters and enhancers which are active in K562 (i.e. overlapping with a GRO-cap TSS and an ENCODE DNase hyper-snesitivity site) by predicted promoters and enhancers is plotted to assess how well models distinguish promoters from enhancers. (E) Predicted enhancers show significantly higher activity than repressed and low coverage regions as measured by a reporter assay (’*’, ‘**’ and ‘***’ indicate p-values <0.05, 0,01 and 0.001). (F) Comparison of experimetal measures of enhancer activity between different studies.

GRO-cap is a very sensitive method that captures also a large amount of TSSs of unstable transcripts. However it is limited to capped RNA species, misses RNAs below the detection threshold and cannot be used to validate repressed (i.e. transcriptionally inactive) regulatory elements. To address these shortcomings we used two additional independent features, TF binding and HOT regions. The binding of TFs to a region of DNA is a pre-requisite for potential regulatory function and transcriptional activity. High Occupancy of Target (HOT) regions are genomic regions which are bound by a large number of different transcription-related factors 12, which were shown to function as enhancers 36 and are enriched in disease-and trait-associated genetic variants 37. As for the benchmark with TSSs, we sorted chromatin states by overlap with HOT regions by decreasing precision and calculated cumulative recall and FDR (Figure 3B). The best performing segmentations for HOT regions were GenoSTAN-Poilog-K562 and GenoSTAN-nb-K562, followed by ChromHMM-ENCODE. The ordering of states with HOT regions was indeed different from the GRO-cap TSSs benchmark. Additionally to GenoSTAN promoter and enhancer states, the repressed enhancer state frequently overlapped with HOT regions with an overall precision of 81% (GenoSTAN-Poilog-K562) and 77% (GenoSTAN-nb-K562). In comparison, the top three ChromHMM-ENCODE states had together a precision of 67%. All other segmentation methods showed a lower precision and recall for HOT regions. This was also reflected in the frequency of individual TF binding sites at enhancer regions, which were generally higher in GenoSTAN enhancer states than in other segmentations (Figure 3C). In particular, only a very small fraction of EpicSeg and Segway-nmeth enhancers were found to be bound by TFs. EpicSeg and Segway-nmeth segmentations were also those with the highest number of predicted enhancers, suggesting that many of these predictions are spurious.

Next, we calculated the recall of FANTOM5 promoters 16 and enhancers 15 to assess how well the models distinguish promoters from enhancers, as it was evident from inspection of specific examples that this distinction was difficult to be established by current methods (Figure 2A). The FANTOM5 consortium have performed extensive mapping of capped transcripts 5’ ends using CAGE and defined enhancers and promoters based on transcriptional activity pattern. FANTOM5 enhancers were defined as regions showing balanced bidirectional capped transcripts, a hallmark of enhancer RNAs 15, whereas FANTOM5 promoters were defined as regions where transcription was biased towards one direction. The FANTOM5 annotation of enhancers and promoters could not entirely replace a chromatin state based approach because (i) the use of expression data in FANTOM5 limits the identified regulatory regions to transcriptionally active elements and (ii) CAGE was shown to be not as sensitive to rapidly degraded transcripts as GRO-cap and therefore might miss regulatory enhancers with unstable transcripts 34. Nonetheless, FANTOM5 provides an annotation of enhancers and promoters based on independent data that is well suited to assess how well the models distinguish promoters from enhancers. We filtered the FANTOM5 annotation to promoters and enhancers for activity in K562 by overlapping them with DHS 11 and GRO-cap TSSs 34. We considered that a promoter state performed well, when the recall of FANTOM5 promoters was high and the recall of FANTOM5 enhancers was low and vice versa for enhancer states. GenoSTAN-Poilog-K562 and ChromHMM-nature enhancer states recall most FANTOM5 enhancers (60%, Figure 3D, Supplementary Table 2). For enhancer states, the recall of FANTOM5 promoters was around 10% except for those of Segway-ENCODE, which recalls almost 35% of FANTOM5 promoters and EpicSeg which 21% FANTOM5 promoters. In accordance with this, many promoter regions were erroneously classified as enhancer regions in this segmentation (e.g. TAL1 promoter in Figure 2A). The recall of FANTOM5 enhancers by promoter states was generally higher (17% - 37%). GenoSTAN-Poilog-K562 and-nb-K562 recalled more than 90% of FANTOM5 promoters and around 20% of FANTOM5 enhancers which is comparable to other studies (Segway-nmeth, ChromHMM-nature, EpicSeg). ChromHMM-ENCODE promoter states had a comparable recall of FANTOM5 promoters (92%), but higher recall of FANTOM5 enhancers (37%) (Figure 3D). This strong overlap of ChromHMM-ENCODE promoters with FANTOM5-labeled enhancers is in accordance with our observation that some enhancer regions were errenuously classified as promoters in ChromHMM-ENCODE (Figure 2A). These results show that GenoSTAN segmentations distinguish promoters from enhancers at similar or better accuracy than other segmentations.

So far we only used indirect evidence (TSSs, HOT regions, TF binding, FANTOM5 enhancer) to draw conclusions about the cis-regulatory activity of a candidate enhancer. As additional and direct evidence for the cis-regulatory activity of enhancer regions inferred by GenoSTAN, we overlapped our enhancers to genomic sequences that were previously tested for cis-regulatory activity in a reporter assay, where candidate elements had been cloned into a plasmid upstream of the promoter of a reporter gene 4. Enhancers from GenoSTAN segmentations showed significantly higher activity than repressed or low coverage regions (GenoSTAN-Poilog-K562 & GenoSTAN-nb-K562: p-value < 0.001 wilcoxon-test, Figure 3E). Interestingly, repressed regions (marked by H3K27me3) showed lower activity than low coverage regions. Moreover, GenoSTAN-Poilog-K562 enhancers showed significantly higher enhancer activity than those of three other studies (Figure 3F), including the original study (p-value < 0.01, ChromHMM-nature enhancers) by Kheradpour et al. 4. This analyis shows that GenoSTAN has higher succes rate in predicting in vivo enhancer activity than previous methods.

#### Comparison of the GenoSTAN, ChromHMM, Segway and EpicSeg algorithms on a common dataset

The K562 genome segmentations of ChromHMM, Segway and EpicSeg used so far were derived from different combinations of data tracks. To verify that the favorable performance of GenoSTAN is mainly due to an improved modeling and not due to different data, we also ran ChromHMM, Segway and EpicSeg on the same data as GenoSTAN-Poilog-K562 and GenoSTAN-nb-K562. Both GenoSTAN-Poilog-K562 and GenoSTAN-nb-K562 had a lower FDR at a similar or higher recall than all three other methods (Supplementary Figure 3). Moreover, we found that changing the binarization for ChromHMM dramatically affected its outcome. Without further manual processing of the data, ChromHMM fitted only one transcriptionally active state, which modeled both promoters and enhancers, regardless of state number (Supplementary Figure 3). We suspected that the high read coverage in the H3K4me1 and H3K4me3 signal tracks made promoters and enhancers indistinguishable after binarization (H3K4me1 and H3K4me3 were called present at both, promoters and enhancers, and they were both called absent elsewhere). When all data tracks were subsampled to the same (and lower) library size, this problem was solved and ChromHMM fitted multiple transcriptionally active states thereby distinguishing promoters from enhancers and, at the same time, increased in accuracy (Supplementary Figure 3). The same problem occurred for Segway, but changing Segway’s parameters did not help distinguish different transcriptionally active chromatin states. To make sure that these results did not depend on the arbitrary choice of the number of states, we ran each method using 10 to 30 states (Methods) and calculated the precision of each state S for recalling HOT (respectively TSS) regions as the fraction of all segments annotated with S that overlapped with a HOT (respectively TSS) region. For each number of states and each segmentation algorithm, we determined the state with highest precision (Supplementary Fig. 4A, B). Independently of the number of states, GenoSTAN-Poilog and GenoSTAN-nb consistently performed best. Even at low state numbers, precision remained constantly high, while it decreased considerably for other methods. We also derived an area under curve (AUC) score for each model, to assess the spatial accuracy in calling TSSs or HOT regions (Supplementary Fig. 4C, D and Methods). Again, AUC scores were consistently highest for the GenoSTAN segmentations.

Altogether this extensive benchmark in the K562 cell line demonstrates that GenoSTAN-Poilog and to a slightly lesser extent GenoSTAN-nb, outperforms current chromatin state annotation algorithms for identifying enhancers and promoters.

### Benchmarks II and III: Chromatin state annotation for ENCODE and Roadmap Epigenomics cell types and tissues

We next applied GenoSTAN to 127 cell types and tissues from ENCODE and Roadmap Epigenomics, the largest compendium of chromatin-related data, using genomic input and the five chromatin marks H3K4me1, H3K4me3, H3K36me3, H3K27me3, and H3K9me3 that have been profiled across the whole compendium 13 (Supplementary Figure 5, Benchmark II, Figure 1B for data tracks and model names). Moreover, we performed a dedicated analysis to 20 of these cell types and tissues which had three further important data tracks: H3K27ac, H3K9ac and DNase-Seq (Supplementary Figure 6, Benchmark III, Figure 1B for data tracks and model names). These further three tracks are important features of active promoters and enhancers, which can lead to more precisely mapped enhancer boundaries 11. We performed similar comparisons as decribed above to the three available segmentations from the Roadmap Epigenomics project with 15, 18 and 25 states (ChromHMM-15, −18, and −25) [13, 38]. All methods were less performant than in Benchmark I, possibly due to lower read coverage or to less rich data. Nonetheless, the GenoSTAN annotations consistently outperformed the existing ones. Specifically, this held when assessing the recovery of FANTOM5 CAGE tags (Figure 4A, assessed for all 127 cell types and tissues), of GRO-cap TSSs (Figure 4B assessed for the cell types with available GRO-Cap TSSs), and of HOT regions (Figure 4C, assessed for the cell types with available HOT regions). Moreover, both GenoSTAN models distinguished better promoters from enhancers than previous annotations (Figure 4D, Supplementary Table 2). The low accuracy of ChromHMM-15 and ChromHMM-18 promoters might be caused by frequent state switching between the promoter and promoter flanking state (Supplementary Figure 7). Consequently, the number of promoter regions in K562 was up to 30% higher in the ChromHMM-15 and −18 segmentations than in the GenoSTAN or ChromHMM-25 segmentations (Supplementary Table 1). The number of predicted enhancers (in K562) also differed greatly. The ChromHMM-15 and −18 state models predict 92,824 (7_Enh) and 22,678 (9_EnhA1) enhancers, while ChromHMM-25 predicts 12,706 and GenoSTAN-Poilog-20 and −127 predict 15,655 and 45,955 enhancers (Supplementary Table 1). Although GenoSTAN predicted more enhancers than the ChromHMM-25 model, the fraction of putative enhancers bound by individual TFs was greater (Figure 4E). For instance 46% (25%) of enhancers were bound by Pol II in the GenoSTAN-Poilog-20 (-127) model, compared to 8%, 18% and 36% in the ChromHMM 15, 18 and 25 state models. Also, the lineage-specific enhancer-binding transcription factor TAL1 binds at 37% (GenoSTAN-Poilog-20) and 27% (GenoSTAN-Poilog-127) of predicted enhancers. Conversely, 13%, 16% and 27% of putative enhancers were bound by TAL1 in the respective 15, 18 and 25 state ChromHMM models (Figure 4E). Collectively, these results show that the improved performance of GenoSTAN is not restricted to the K562 dataset.

**Figure 4:**
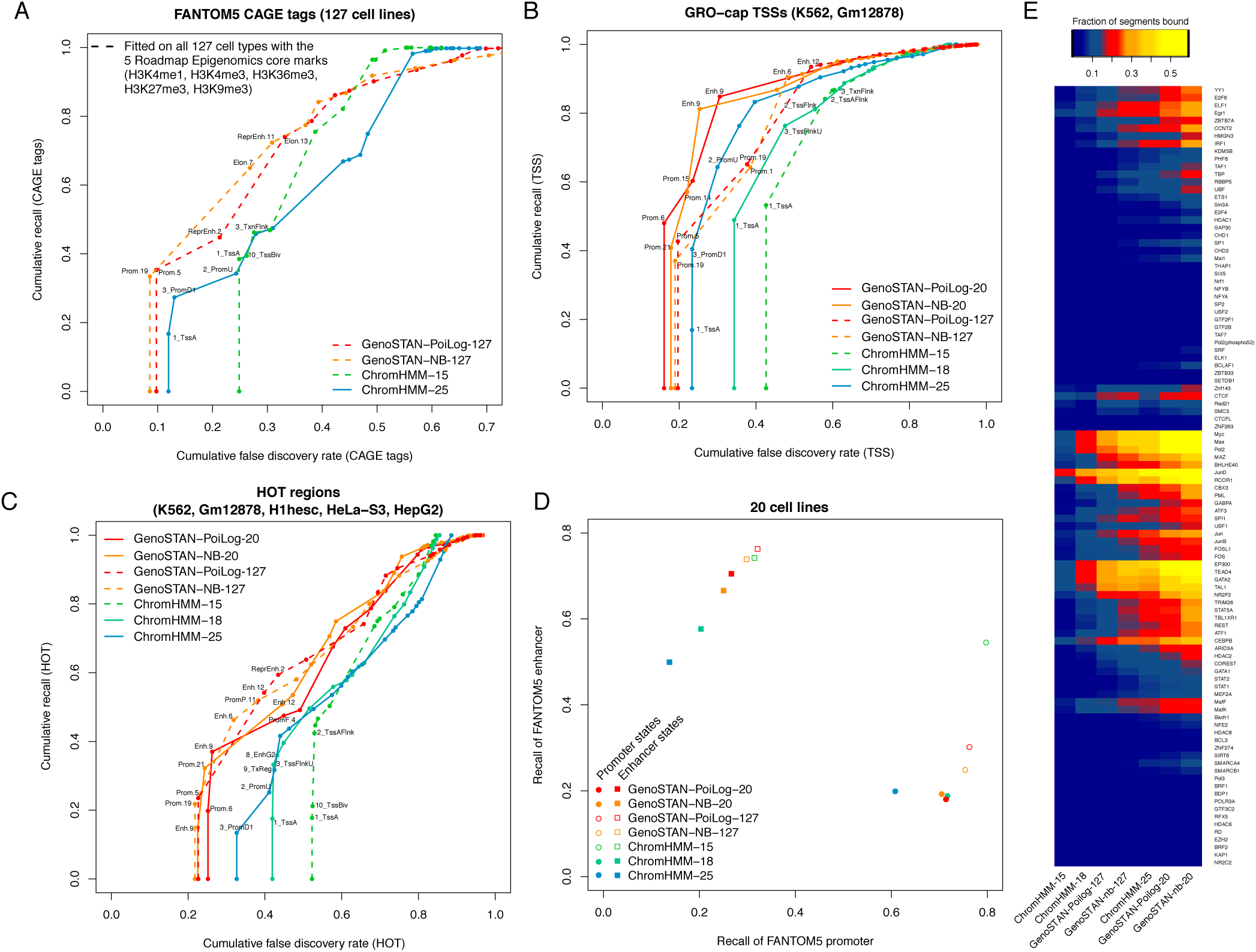
Comparison of GenoSTAN to other published segmentations on benchmark II and III. (A) Performance of chromatin states in recovering FANTOM5 CAGE tags in 127 cell types. CAGE tags were verlapped with chromatin states wihout the use of cell type information. Cumulative FDR and recall are calculated by subsequently adding states (in order of increasing FDR). (B) Performance of chromatin states in recovering GRO-cap transcription start sites in two cell types where GRO-cap data was available. (C) The same as in (B) for ENCODE HOT regions for five cell types where annotation of HOT regions was available. (D) Recall of FANTOM5 promoters and enhancers by predicted promoters and enhancersis plotted to assess how well models distinguish promoters from enhancers. (E) The fraction of predicted enhancer segments bound by individual TFs is shown for different studies. GenoSTAN enhancers are more frequently bound by TFs than those from other studies.

### Cell-type specific enrichment of disease-and other complex trait-associated genetic variants at promoters and enhancers

Previous studies showed that disease-associated genetic variants are enriched in potential regulatory regions [13, 21, 39, 40, 41, 42] demonstrating the need for accurate maps of these elements to understand genotype-phenotype relationships and genetic disease. To study the potential impact of variants in regulatory regions on various traits and diseases, we overlapped our enhancer and promoter annotations from 127 cell types and tissues (Benchmark II, Figure 1B) with phenotype-associated genetic variants from the NHGRI genome-wide assocation studies catalog (NHGRI GWAS Catalog 43). First, we intersected trait-associated variants with enhancer and promoter states (GenoSTAN-Poilog-127). Overall, 37% of all trait-associated SNPs were located in potential enhancers and 7% in potential promoters. The number of traits significantly enriched (at FDR <0.05) with enhancers or promoters in at least one cell type or tissue was larger for GenoSTAN-Poilog-127 (69 traits for GenoSTAN-Poilog-127 for enhancers and 20 traits for promoters) than for the best performing ChromHMM-model (ChromHMM-15, 64 traits for enhancers and 18 traits for promoters). The better performance of GenoSTAN-Poilog-127 was found at all FDR cutoffs (Supplementary Figure 8). To control for the fact that methods can differ among each other regarding the length of the promoters and enhancers they predict, we furthermore computed the recalls of GWAS variants for a fixed genomic coverage. Restricting to a total genomic coverage of 2% (random subsetting, also allowing confidence interval computation, Methods), enhancers of all GenoSTAN models overlaped a higher fraction of GWAS variants at a similar to better per base pair density compared to the current ChromHMM annotations (Figure 5A). The same trend was observed for promoters when restricting to 1% of genomic coverage (Figure 5B). The improved overlap with trait-associated variants indicates that GenoSTAN annotation has a higher enrichment for functional elements than the current annotation.

**Figure 5:**
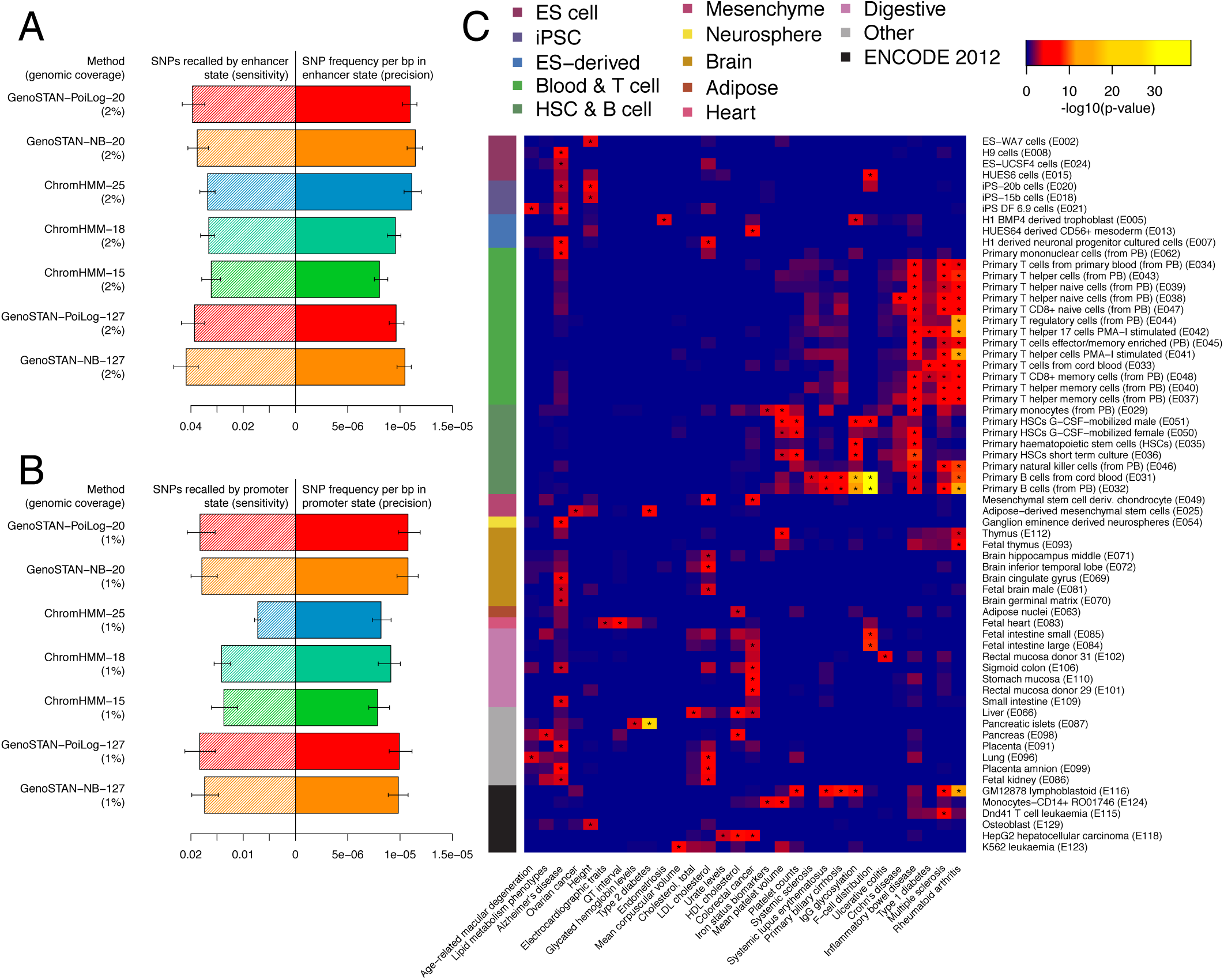
Enrichments of genetic variants associated with diverse traits in enhancers and promoters are specific to the relevant cell types or tissues. (A) Median SNP recall and frequency was calculated for enhancer states in different segmentations by restricting it to a total genomic coverage of 2% (100 samples of random subsetting) to control for different number of enhancer calls between the segmentations. Error bars show the 95% confidence interval. (B) The same as in (A) but for promoters. (C) The heatmap shows the-log10(p-value) of significantly enriched traits in enhancer states (GenoSTAN-Poilog-127, p-value < 0.001, marked by ‘*’). Only cell types and tissues where at least one trait was significantly enriched are shown. P-values were adjusted for multiple testing using the Benjamin-Hochberg correction.

In accordance with previous studies [13, 21] we found that individual variants were strongly enriched in enhancer or promoter states specifically active in the relevant cell types or tissues (Figure 5C, Supplementary Figure 8C). Variants associated with height were significantly associated with osteoblasts (at FDR <0.001 here and after, performed on Benchmark II for consistency across cell types and tissues). Variants associated with immune response or autoimmune disorders were enriched in B-and T-cell enhancers (Figure 5C) and promoters (Supplementary Figure 8C). These incude for instance HIV-1 control, autoimmune disease associated SNPs for systemic lupus erythematosus, inflammatory bowel disease, Ulcerative colitis, Rheumatoid arthritis, Primary biliary cirrhosis and Multiple sclerosis. Variants associated with electrocardiographic traits and QT interval were enriched in fetal heart enhancers. SNPs associated with colorectal cancer were enriched in enhancers specific to the digestive system. These results illustrate that the annotation of potential promoters and enhancers generated in this study can be of great use for interpreting genetic variants associated, and underscore the importance of cell-type or tissue specific annotations.

## A novel annotation of enhancers and promoters in human cell types and tissues

We then compiled the results from the best performing annotations for each cell type and tissue into a single annotation file. The combined annotation file is available as Supplementary Data. All individual chromatin state annotations are available at i12g-gagneurweb.in.tum.de/public/paper/GenoSTAN. For the combined annotation file, we chose GenoSTAN with Poisson-lognormal in every instance, as it performed best in almost every comparison we conducted. We used the results from benchmark I for K562, from benchmark III for the 20 cell types and tissues, and from benchmark II for all the remaining Roadmap Epigenomics cell types and tissues. Overall, our annotation reports typically between 8,945 and 16,750 (10% and 90% quantiles of number of promoters across all 127 cell types and tissues) active promoters per cell type or tissue. This number is consistent with the typical number of expressed genes per tissue (in 11,953 to 16,869 range, 44). However, the median width of these elements depends on the data on which the annation was based. For the benchmark III dataset, promoters are much narrower (800bp median) than for the K562 annotations (1.4 kb, Benchmark I data sst), suggesting that promoter regions in the 20 cell types more accurately recover DNase hypersensitivity sites (DHS) of the core promoter (Figure 2, Supplementary Figure 6). The number of enhancers per cell type or tissue varied more greatly (between 8,208 and 33,596 for the 10% and 90% quantiles). The large variation of the number of enhancers might be partly due to differences of sensitivity in complex biological samples. Consistent with this hypothesis, much fewer enhancers were identified in tissues than in primary cells and cell lines (Supplementary Figure 9) likely because enhancers that are active only in a small subsets of all cell types present of a tissue may be not detected. As more cell-type specific data will be available, improved maps can be generated. The GenoSTAN software, which is publicly available, will be instrumental to update these genomic annotations.

## Promoters and enhancers have a distinct TF regulatory landscape

The biochemical distinction between enhancers and promoters is a topic of debate [6, 7]. We explored to which extent enhancers and promoters are differentially bound by TFs using the K562 cell line dataset because i) we obtained the most accurate annotation for this cell line (GenoSTAN-Poilog-K562, Benchmark I) and ii) ChIP-seq data was available for as many as 101 TFs in this cell line 11. Nine TF modules were defined by clustering based on binding pattern similarity across enhancers and promoters (Methods, Figure 6). These 9 TF modules were further characterized by the propensity of their TFs to bind promoters, enhancers or both (Figure 6). In accordance with previous studies [45, 46], this recovered many complexes and promoter-associated and enhancer-associated proteins, including the CTCF/cohesin complex (CTCF, Rad21, SMC3, Znf143), the AP-1 complex (Jun, JunB, FOSL1, FOS), Pol3, promoter and enhancer associated modules, and factors associated with chromatin repression (EZH2, HDAC6).

**Figure 6:**
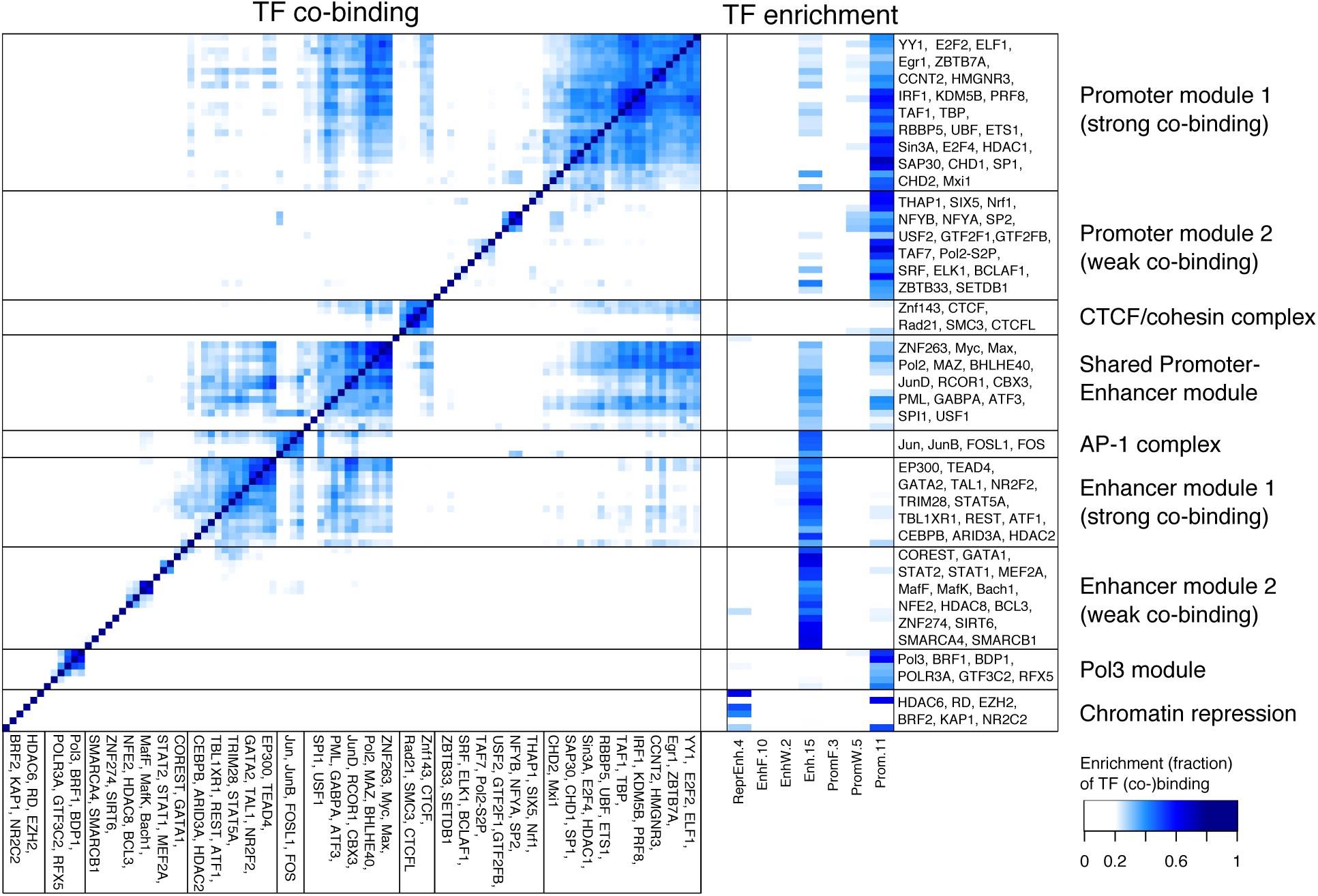
Promoters and enhancers have a distinctive TF regulatory landscape. Co-binding (left) and enrichment of transcription factor binding sites (right) in chromatin states (GenoSTAN-Poilog-K562) for 101 transcription factors in K562 reveals TF regulatory modules with distinct binding preferences for promoters, enhancers and repressed regions. The co-binding is depicted as the frequency of binding sites of two TFs that co-occur in a chromatin state divided by the number of all binding sites of the two TFs (Jaccard index). For each TF, enrichments were normalized to sum up to 1 across all 18 chromatin states of GenoSTAN-Poilog-K562.

Moreover, the modules identified provided insights into the distinction of promoters and enhancers. On the one hand, some TFs are common to both enhancers and promoters, which supports previous reports [15, 7]. In accordance with the recent finding of widespread transcription at enhancers 34, Pol II and multifunctional TFs Myc, Max, and MAZ 47 are part of a TF module - which we called the Promoter-Enhancer-Module (PEM) - which had approximately equal binding preferences for promoter and enhancer states, but also co-localized with other TFs specifically binding enhancers or promoters (Figure 6).

On the other hand enhancers and promoters were also bound by distinct TFs, which is consistent with previously reported TF co-occurrence patterns at gene-proximal and gene-distal sites [46, 45]. Among the promoter and enhancer-associated proteins we defined Promoter module 1 and 2 (PM1, PM2), Enhancer module 1 and 2 (EM1, EM2), which had a strong preference for binding either a promoter or an enhancer, but exhibited different co-binding rates (Figure 6). Promoter module 1 contained TFs which were specifically enriched in promoter states and associated with basic promoter functions, such as chromatin remodeling (CHD1, CHD2), transcription initiation or elongation (TBP, TAF1, CCNT2, SP1) and other TFs involved in the regulation of specific gene classes (e.g. cell cycle: E2F4) 47. However, it also included TFs known as transcriptional repressors (e.g. Mxi1, a potential tumor suppressor, which negatively regulates Myc). While TFs in PM1 showed a high co-binding rate, PM2 factors exhibited low co-binding. This might be partially explained by lower efficiency of the ChIP, since PM2 also contained general TFs such as TFIIB, TFIIF or the Serine 2 phospho-isoform of Pol II, which are expected to co-localize with other general TFs from PM1.

EM1 contained TFs with high co-binding rate, which included TAL1, an important lineage-specific regulator for erythroid development (K562 are erythroleukemia cells) and which had been shown to interact with CEBPB, GATA1 and GATA2 at gene-distal loci [46, 48]. It also contained the enhancer-specific transcription factor P300 32 and transcriptional activators (e.g. ATF1) and repressors (e.g. HDAC2, REST) 47. Analogously to PM2, EM2 contained enhancer-specific transcriptional activators and repressors with a low co-binding rate.

Altogether this analysis highlights the common and distinctive TF binding properties of enhancers and promoters.

## Discussion

We introduced GenoSTAN, a method for *de novo* and unbiased inference of chromatin states from genome-wide profiling data. In contrast to previously described methods for chromatin state annotation, GenoSTAN directly models read counts, thus avoiding data transformation and the manual tuning of thresholds (as in ChromHMM and Segway), and variance is not shared between data tracks or states (as in EpicSeg and Segway) [28, 8, 9]. GenoSTAN is released as part of the open-source R/Bioconductor package STAN [29, 49, 50], which provides a fast, multiprocessing implementation that can process data from 127 human cell types in less 3-6 days (GenoSTAN-Poilog-127: 6 days,-nb: 3 days).

Application of GenoSTAN significantly improved chromatin state maps of 127 cell types and tissues from the ENCODE and Roadmap Epigenomics projects [11, 13]. Binding of enhancer-associated co-activator CBP and histone acetyltransferase P300 was used by several studies for the genome-wide prediction of enhancers [32, 31, 27]. From these predictions a distinctive chromatin signature for promoters and enhancers was derived based on H3K4me1 and H3K4me3 27. In particular, the ratio H3K4me1/H3K4me3 was found to be low at promoters, in comparison to enhancers. Active and poised enhancers could also be distinguished by presence or absence of H3K27me3 and H3K9me3 51. All these features could be confirmed by GenoSTAN, making it a promising tool for the biochemical characterization of enhancers and promoters. Moreover, extensive benchmarks based on independent data including transcriptional activity, TF binding, cis-regulatory activity, and enrichment for complex trait-associated variants showed the highest accuracy of GenoSTAN annotations over former genome segmentation methods.

The GenoSTAN annotation sheds light on the common and distinctive features of promoters and enhancers, which currently are an intense subject of debate [6, 7]. Among other characteristics, a shared architecture of promoters and enhancers was proposed based on the recent discovery of widespread bidirectional transcription at enhancers [7, 34, 35]. This was supported by the observation that enhancers, which are depleted in CpG islands have similar transcription factor (TF) motif enrichments as CpG poor promoters 15. However, another study showed that TF co-occurrence differed between gene-proximal and gene-distal sites [46, 45]. GenoSTAN chromatin states revealed a very distinct TF regulatory landscape of these elements and therefore suggest that promoters and enhancers are fundamentally different regulatory elements, both sharing the binding of the core transcriptional machinery. Our annotation of enhancers and promoters will be a valuable resource to help characterizing the genomic context of the binding of further TFs.

Indirectly, our analysis showed that chromatin state annotations are better predictors of enhancers than the transcription-based definition provided by the FANTOM5 consortium 15. While FANTOM5 enhancers are an accurate predictor for transcriptionally active enhancers, the sensitivity remains poor (only 4,263 enhancers were called by overlap with GRO-cap TSSs and DHS, which is less than the estimated number of transcribed genes, for K562 cells compared to about 20,000-30,000 for ChromHMM and 10,000-20,000 for GenoSTAN). Although, the sensitivity of the transcription-based approach can increase with transient transcriptome profiling [52, 53] or nascent transcriptome profiling 54, the chromatin state data undoubtedly add valuable information for the indentification of promoters and enhancers. Because it models count data, GenoSTAN analysis can in principle also integrate RNA-seq profiles, for instance using it in a strand-specific fashion 29.

Systematic identification of cis-regulatory active elements by direct activity assays is notoriously difficult. STARR-Seq for instance is a high-throughput reporter assay for the *de novo* identification of enhancers 5. It was previously used to identify thousands of cell-type specific enhancers in *Drosophila*, but has not been applied to human yet. Moreover, STARR-Seq makes rigid assumptions about the location of the enhancer element with respect to the promoter, and it does not account for the native chromatin structure. This might identify regions that are inactive *in situ* 5. Other experimental assays for the validation of predicted ENCODE enhancers lead to different results [3, 4]. Complementary to these approaches, the systematic evaluation of cis-regulatory activity based on candidate regions in human cells have made progress with the advent of high-throughput CRISPR perturbation assays 55. Because it requires candidate cis-regulatory regions in a first place, such approach will benefit from improved annotation maps as the one we are providing.

Thus, we foresee GenoSTAN to be instrumental in future efforts to generate robust, genome-wide maps of functional genomic regions like promoters and enhancers

## Methods

### Availability of GenoSTAN and chromatin state annotations

GenoSTAN is freely available from bioconductor.org/ as part of our previously published R/Bioconductor package STAN 29. Chromatin state annotations for benchmark I, II and III can be downloaded from i12g-gagneurweb.in.tum.de/public/paper/GenoSTAN. The combined promoter and enhancer annotation is available as Supplementary Data.

### Motivation of Poisson-lognormal and negative binomial emissions

The Poisson-lognormal and the negative binomial distribution can be thought of as extensions of the Poisson distribution that allow for greater variance. We will now motivate both distributions from a Poisson distribution with a prior on the mean of the Poisson.

Suppose that *X* ∼ *Poisson*(*x*|Λ) is a Poisson random variable and Λ ∼ *Gamma* (λ|*α,β*). From this we can derive the negative binomial with success rate *p* and size *r* 56:

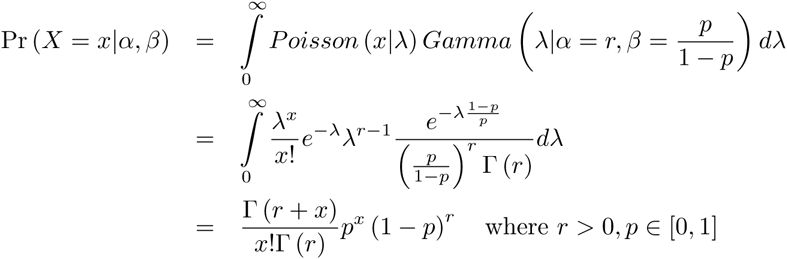

In order to increase interpretability in the context of read counts, we re-parameterize this with mean 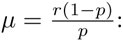:

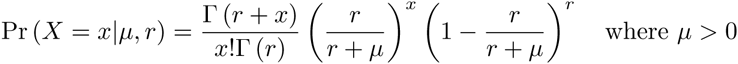

The Poisson-lognormal distribution can be motivated likewise. Assume that *X* ∼ *Poisson*(*x*|Λ) is a Poisson random variable and Λ ∼ *N* (log (λ) |*μ, σ*). Then the Poisson-lognormal is given by 57:

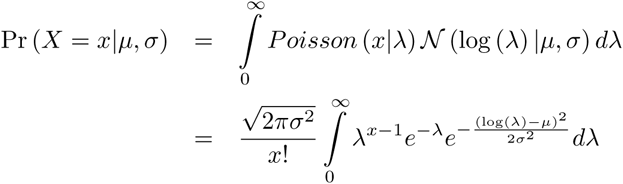

A closed form solution for this distribution does not exist. Thus numerical integration is needed to calculate probabilities, which is done in GenoSTAN by using the R package poilog [58, 49].

### Optimization of Poisson-lognormal and negative binomial emissions

Let 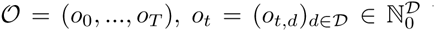 be an obsevational sequence of |𝔇|-dimensional count vectors *o*_*t*_. An HMM assumes that each observation *o*_*t*_ is *emitted* by a corresponding hidden (unobserved) variable *s*_*t*_, *t* = 0, …,*T*. A hidden variable can assume values from a finite set of states *𝒦*. Each state *𝒦* ∈ 𝒦 is associated to an emission distribution *ψ*_*k*_, which defines the probability of making a certain observation, *ψ*_*k*_(o_t_). GenoSTAN assumes that the components *o*_*t,d*_, d ∈ 𝔇, of a single observation *o*_*t*_ are independent, and hence 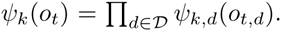. The value of *s*_*t*_ determines the probability of observing *o*_*t*_ by Pr(*o*_*t*_ | *s*_*t*_) = *ψ*_*st*_(*o_t_*). HMM learning is carried out using the Baum-Welch algorithm 25. The optimization problem for the paramters of a single emission distribution *ψ*_*i,d*_ can be written as

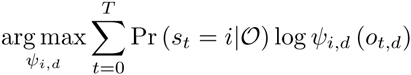

where Pr(*s*_*t*_ = *i* | *𝒪*) is calculated efficiently by the Forward-Backward algorithm, and *ψ*_*i,d*_ is maximized within the class of negative binomial or Poisson-lognormal distributions. An analytical solution for this problem does not exist. Thus, we resort to numerical optimzation. As indicated by 28, above formula can be very costly to compute, since the function needs to evaluate a sum over the complete observation sequence (i.e. the complete binned genome) in each iteration. However, computations are greatly simplified by grouping together observations *o*_*t,d*_ with the same count number. Let *𝒞*_*d*_ be the set of unique counts in dimension *d*. Then the following terms can be precomputed for all *c* ∈ *𝒞*_*d*_ before optimization:

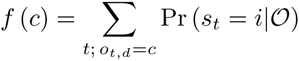

The objective function becomes

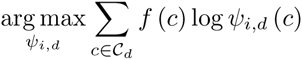

which avoids redundant calculations of *ψ*_*i,d*_(*o*_*t*_), *t* = 0,…,*T*, and greatly reduces complexity since |𝒞_*d*_| ≪ *T*.

### Correction for library size

The sequencing depth can be very different between experiments. GenoSTAN addresses this problem by using pre-computed scaling factors to correct for varying sequencing depths for a data track between cell types. In this work, the ‘total count’ method is used 59. Let *𝓛* be the set of cell types and *r*_*d,l*_ the number of reads of data track *d* ∈ 𝔇 in cell line *l* ∈ *𝓛*. The scaling factor is then computed as

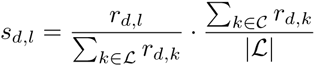

The probability of an observation *o*_*t,l*_ is calculated as 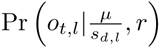 in the case of negative binomial and 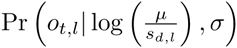 in the case of Poisson-lognormal emissions.

### Model initialization

Initialization of model parameters is crucial for HMMs since the EM algorithm is a gradient method which converges to a local maximum. K-means is a widely used approach to derive an initial clustering to estimate model parameters 25. In order to make this approach applicable to sequencing data, we added a pseudocount and log-transformed the data before k-means clustering. However, without further processing k-means rarely converged and the procedure was slow on the complete data set. To address these issues, we further processed and filtered the data. First, a threshold for signal enrichment for each data track is calculated using the default binarization approach of ChromHMM 8. The threshold is the smallest discrete number *nd* > 0 such that Pr (*X* > *n*_*d*_) < 10^−4^ where *X* is a Poisson random variable with mean 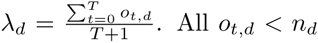 were set to 0, which improved convergence of k-means. To improve the speed, all genomic bins *o*_*t,d*_ where ∀*d* ∈ 𝔇 : *o*_*t,d*_ = 0 were removed and defined as a ‘background cluster’. K-means was then run on the rest of the data with *|𝒦|* − 1 clusters. This clustering (the ‘background’ and k-means clusters) was then used to derive an initial estimate of emission function parameters. Initial state and transition probabilities were initialized uniform.

### Data preprocessing

Benchmark I (K562 ENCODE) sequencing data was mapped to the hg20/hg38 (GRCh38) genome assembly (Human Genome Reference Consortium) using Bowtie 2.1.0 60. Samtools 61 was used to quality filter SAM files, whereby alignments with MAPQ smaller than 7 (-q 7) were skipped. To obtain midpoint positions of the ChIP-Seq fragments, the (single end) reads were shifted in the appropriate direction by half the average fragment length as estimated by strand coverage cross-correlation using the R/Bioconductor package chipseq 50. Next, ChIP-Seq tracks were summarized by the number of fragment midpoints in consecutive bins of 200 bp width. The data for the 127 ENCODE and Roadmap Epigenomics cell types (benchmark II and III) was downloaded as preprocessed tagAlign files from the Roadmap Epigenomics supplementary website 13. Fragment length was again estimated using the R/Bioconductor package chipseq and reads were shifted by the fragment half size to the average fragment midpoint 50. The genome was partitioned into 200bp bins and reads were counted within each bin.

### Model fitting of GenoSTAN

GenoSTAN was fitted on the complete data of benchmark data set I. The signal used for GenoSTAN model training on Benchmark data set II and III was extracted from ENCODE pilot regions (1% of the human genome analyzed in the ENCODE pilot phase 62) for each cell type, which together covered 20% and 127% of the human genome. The GenoSTAN-nb-20 model was learned in one day, the GenoSTAN-Poilog-20 model in two days using 10 cores. Model learning on Benchmark set II using 10 cores took three (GenoSTAN-nb-127) and six days (GenoSTAN-Poilog-127). Precomputed library size factors were used to correct for variation in read coverage.

### Model fitting of ChromHMM, Segway and EpicSeg

The data was binarized as described in 8 and ChromHMM was fitted with default parameters. Before applying Segway, the data was transformed using the hyperbolic sine function 9 and a runing mean over a 1kb sliding window was computed to smooth the data. Segway was fitted on ENCODE pilot regions using a 200bp resolution. EpicSeg was fitted on the untransformed count data with default parameters.

### Processing of chromatin state annotations and external data

All state annotations and external data were lifted to the hg20/hg38 (GRCh38) genome assembly using the liftOver function from the R/Bioconductor package rtracklayer 63. Overlap of state annotations with external data was calculated with GenomicRanges 64.

### Computation of area under curve

AUC values were calculated on Benchmark set I for GenoSTAN, ChromHMM, Segway and EpicSeg. To this end, a segmentation was transformed into a binary classifier and evaluated as follows. Each 200bp bin in the genome overlapping with HOT (TSSs) regions was considered as ‘true condition’, the rest as ‘false’. For each state S the precision for recalling HOT (TSS) regions was calculated as the fraction of all segments annotated with S that overlapped with a HOT (TSS) region. States were then sorted by decreasing precision. The rank of each state was used as score in the prediction of HOT (TSS) regions on each 200bp bin in the genome, which was then used to calculate AUC values.

### Analysis of transcription factor (co-)binding

TF enrichment in chromatin states was calculated as described earlier 45. Let *TF*^*nt*^ be the total number of nucleotides in the binding sites (peaks) a TF and 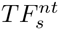 the number of nucleotides in the binding sites that overlap with state *s*. Further let *s*^*nt*^ be the total number of nucleotides in the genome covered by state s and let *l* be the length of the genome. TF enrichment is then calculated as 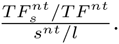. For each TF, enrichments were normalized to sum up to 1 across all 18 chromatin states (GenoSTAN-Poilog-K562). The co-binding rate was calculated as the frequency of binding sites of two TFs that co-occur in a chromatin state divided by the number of all binding sites of the two TFs (Jaccard index).

### Tissue-specific enrichment of disease-and complex trait-associated variants in regulatory regions

The GWAS catalog was obtained from the gwascat package from Bioconductor [50, 43]. Statistical testing was carried out in a similar manner as described in 13. The enrichment of SNPs from individual genome-wide association studies was calculated for traits with at least 20 variants. SNPs for each trait were overlapped with promoter and enhancer regions and tested against the rest of the GWAS catalogue as background using Fisher’s exact test. P-values were adjusted for multiple testing using the Benjamini-Hochberg correction. In order to calculate the recall and frequency of SNPs, promoter and enhancer states were randomly sampled until a genomic coverage of 2% for enhancers and 1% of promoters was reached. This was done to control for the fact that methods can differ among each other regarding the length of the promoters and enhancers they predict. This procedure was repeated 100 times enabling the calculation of 95% confidence intervals.

## Acknowledgments

We thank Lars Steinmetz, Judith Zaugg, Aino Jarvelin, Wu Wei and Philip Brennecke for fruitful discussions and hosting BZ during the early phase of the project. BZ was supported by a DAAD short term research grant.

## Author contributions

BZ, JG and AT developed the statistical methods and computational workflow of the study. BZ developed and implemented all software and scripts and carried out all computational analyses. BS helped with preprocessing of the K562 data set. MM and PC helped with interpretation of the biological results. BZ, JG and AT wrote the manuscript with input from all authors. All authors read and approved the final version of the manuscript.

## Conflict of interest

The authors declare that they have no conflict of interest.

## Supplementary Figures

**Supplementary Figure 1:**
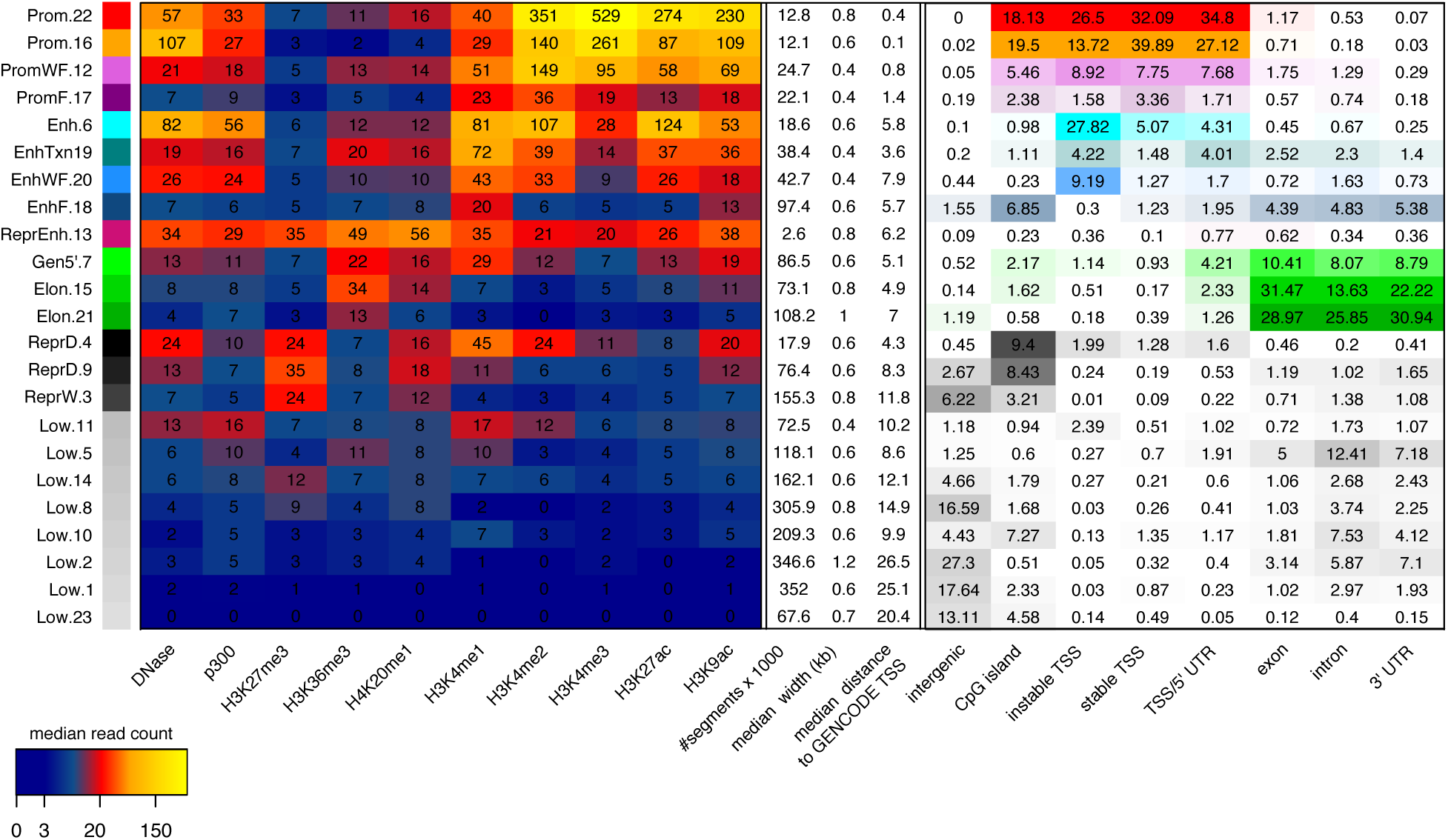
Median read coverage of GenoSTAN-nb-K562 chromatin states (left), their number of annotated segments in the genome, their median width and distance to the closest GENCODE TSS (middle). The right panel shows recall of genomic regions by chromatin states.

**Supplementary Figure 2:**
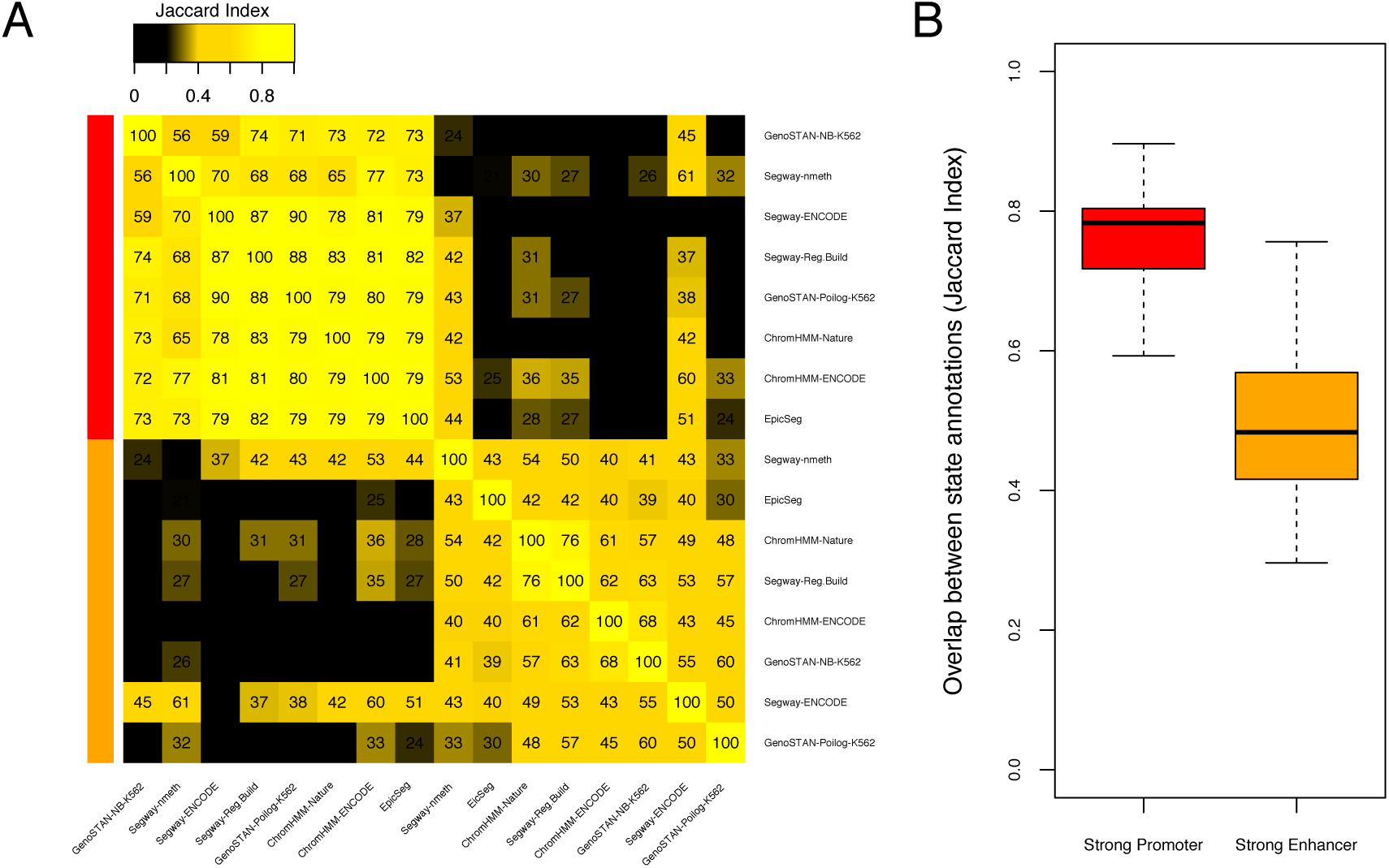
(A) Heatmap of pairwise overlap (Jaccard index) of promoter (red) and enhancer (orange) state annotations from different studies on benchmark I. Rows and columns were ordered by separate clustering of promoter and enhancer overlaps. (B) Distribution of pairwise Jaccard indices for strong promoters and enhancers (off-diagonal elements of promoter and enhancer sub-matrices from (A)).

**Supplementary Figure 3:**
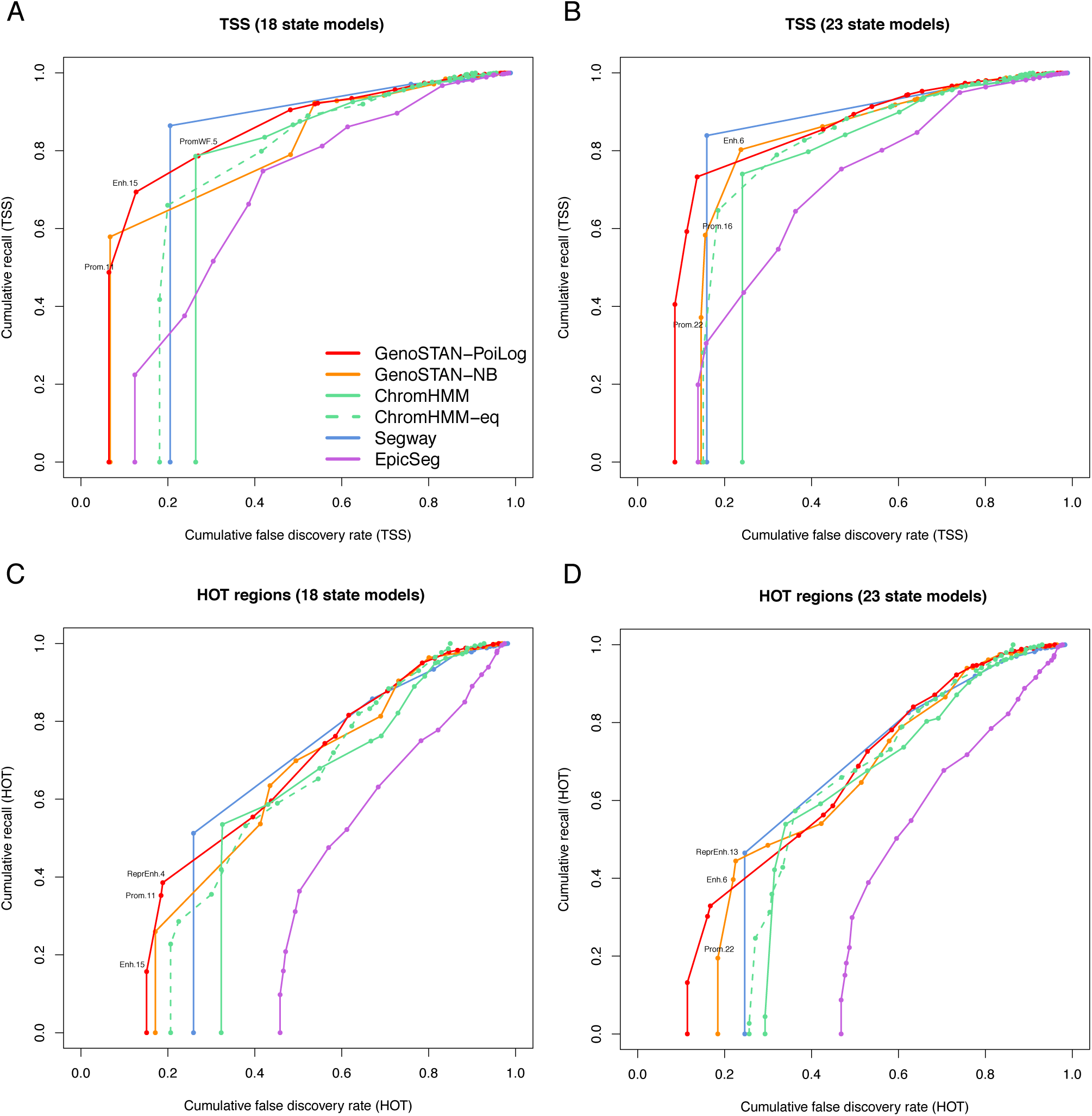
Comparison of GenoSTAN to other methods using 18 and 23 states on the same data set in K562 (Benchmark I). (A-B) Performance of chromatin states in recovering GRO-cap transcription start sites for the 18 and 23 state models. Cumulative FDR and recall are calculated by subsequently adding states (sorted by decreasing precision). (C-D) The same as in (A-B) for ENCODE HOT regions.

**Supplementary Figure 4:**
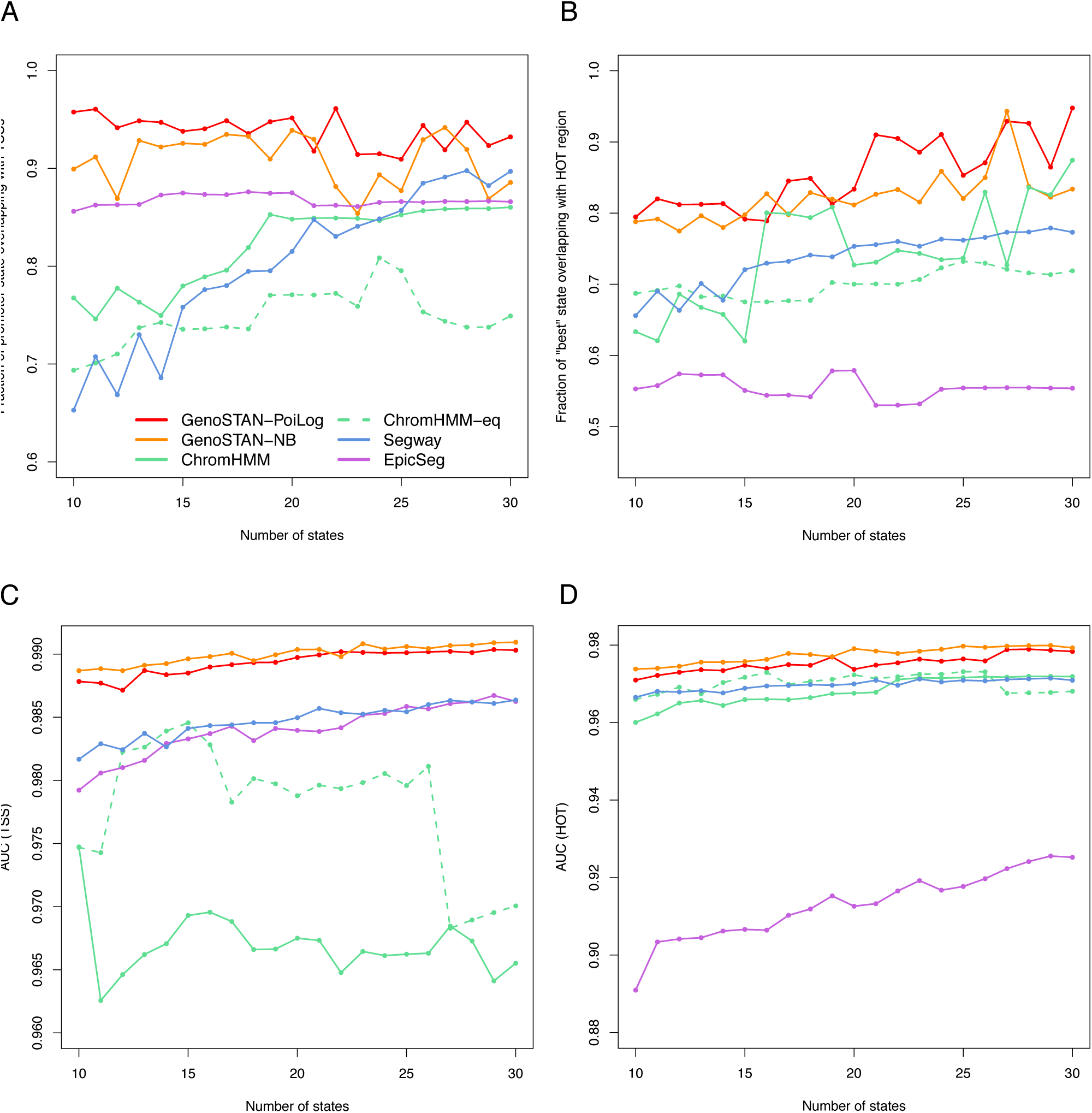
Comparison of chromatin segmentation algorithms with respect to their ability to call GRO-cap transcription start sites (left panels) and ENCODE HOT regions (right panels), as a function of the state number used in the respective algorithm (x-axes). All models were learned on the data set of benchmark I. (A-B) For each model, the state with highest precision in recalling HOT (respectively TSS) regions is shown. (C-D) For each model, an area under curve (AUC) score (see Methods) is plotted to asses the spatial accuracy of a genome segmentation.

**Supplementary Figure 5:**
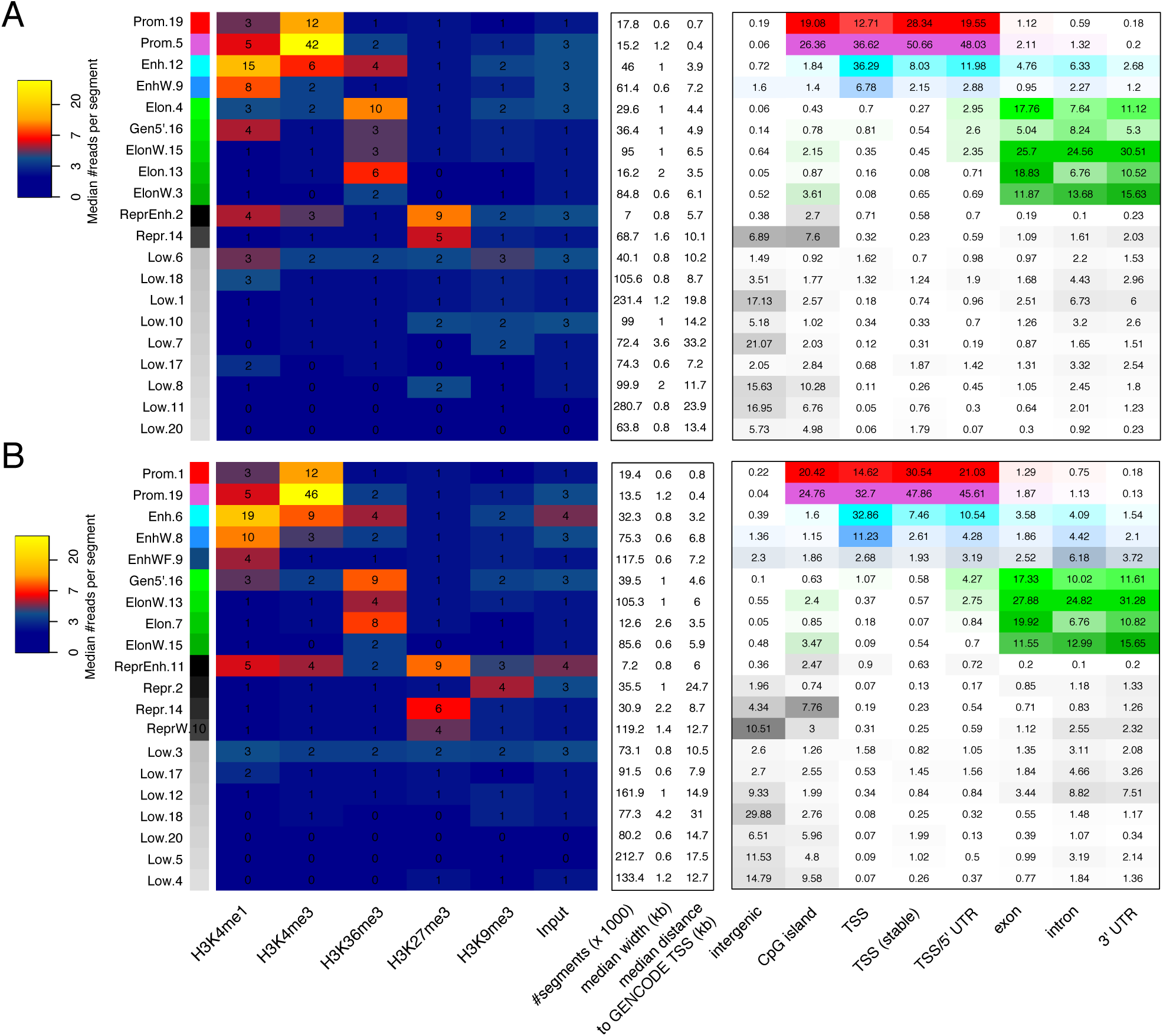
GenoSTAN models for benchmark II. (A) Median read coverage of GenoSTAN-Poilog-127 (Benchmark set II, fitted on the 127 ENCODE and Roadmap Epigenomics cell types and tissues) chromatin states (left), their number of annotated segments in the genome, their median width and distance to the closest GENCODE TSSs of segments (middle). The right panel shows recall of genomic regions by chromatin states. (B) The same as (A) for GenoSTAN-nb-127.

**Supplementary Figure 6:**
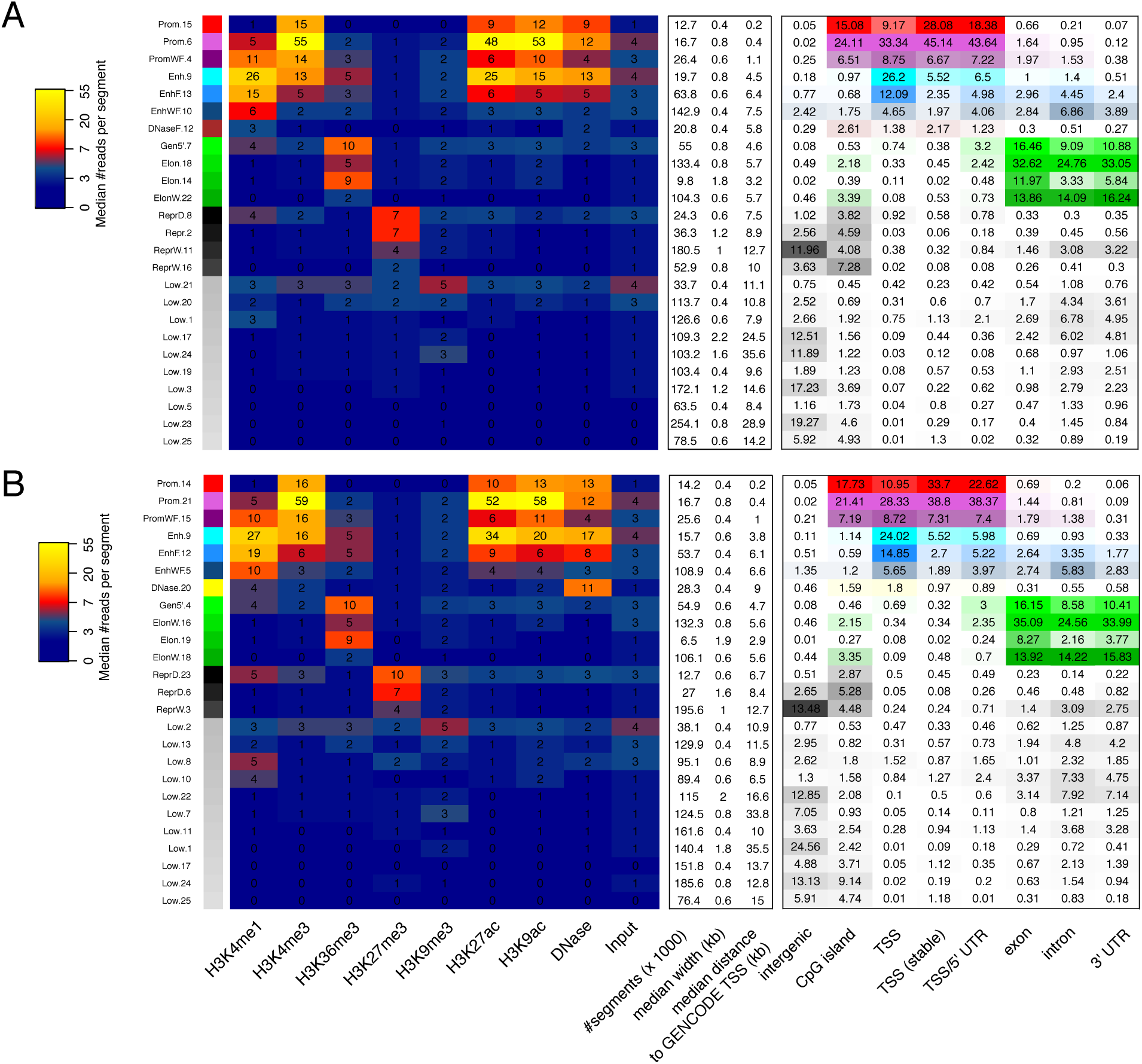
GenoSTAN models for benchmark III. (A) Median read coverage of GenoSTAN-Poilog-20 (Benchmark set III, fitted on the 20 ENCODE and Roadmap Epigenomics cell types and tissues) chromatin states (left), their number of annotated segments in the genome, their median width and distance to the closest GENCODE TSSs of segments (middle). The right panel shows recall of genomic regions by chromatin states. (B) The same as (A) for GenoSTAN-nb-20.

**Supplementary Figure 7:**
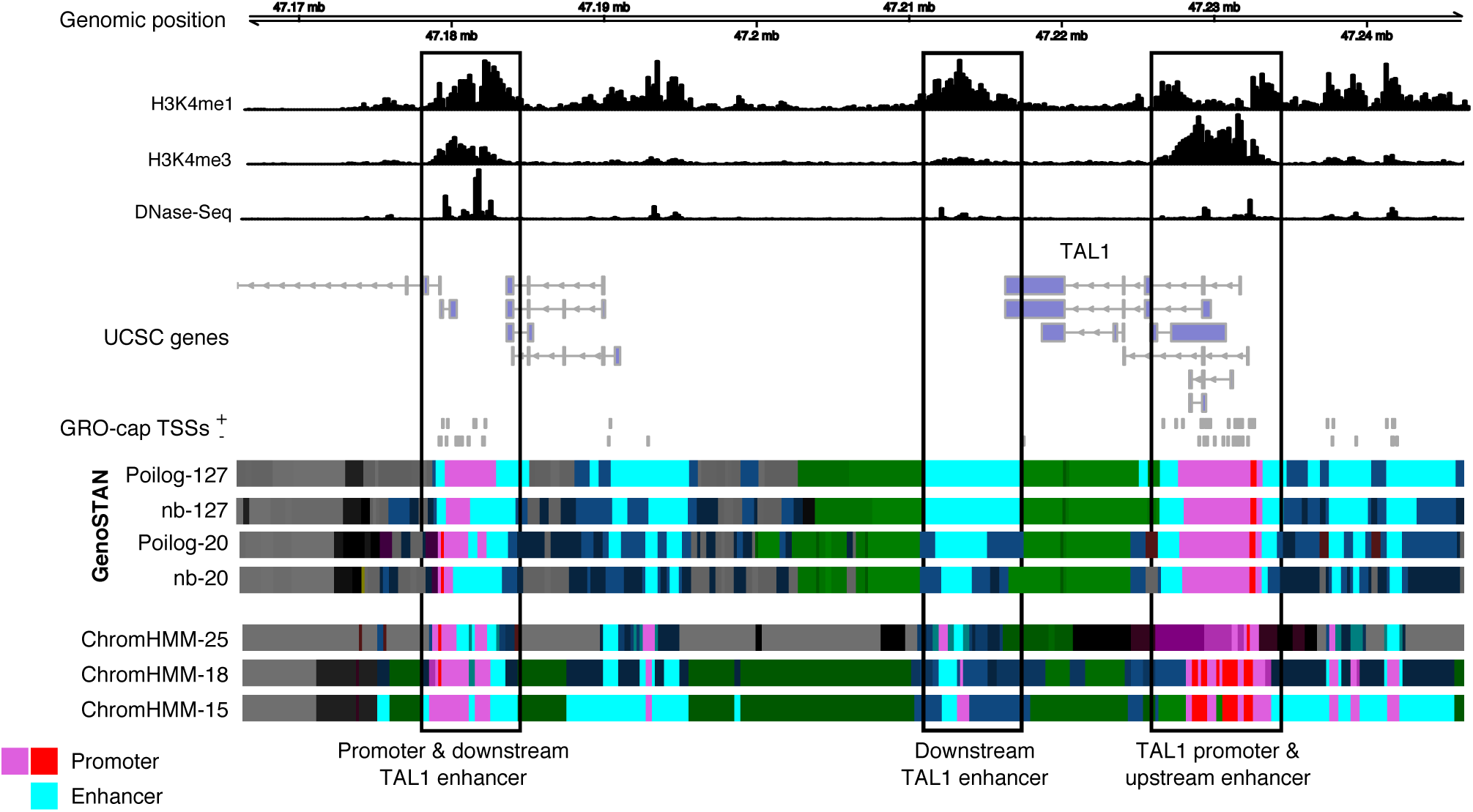
Chromatin states annotations for benchmark II and III GenoSTAN segmentations are shown with the three Roadmap Epigenomics ChromHMM segmentations with 15 (ChromHMM-15), 18 (ChromHMM-18) and 25 (ChromHMM-25) states. Shown is the TAL1 locus with segmentations and data from the K562 cell line.

**Supplementary Figure 8:**
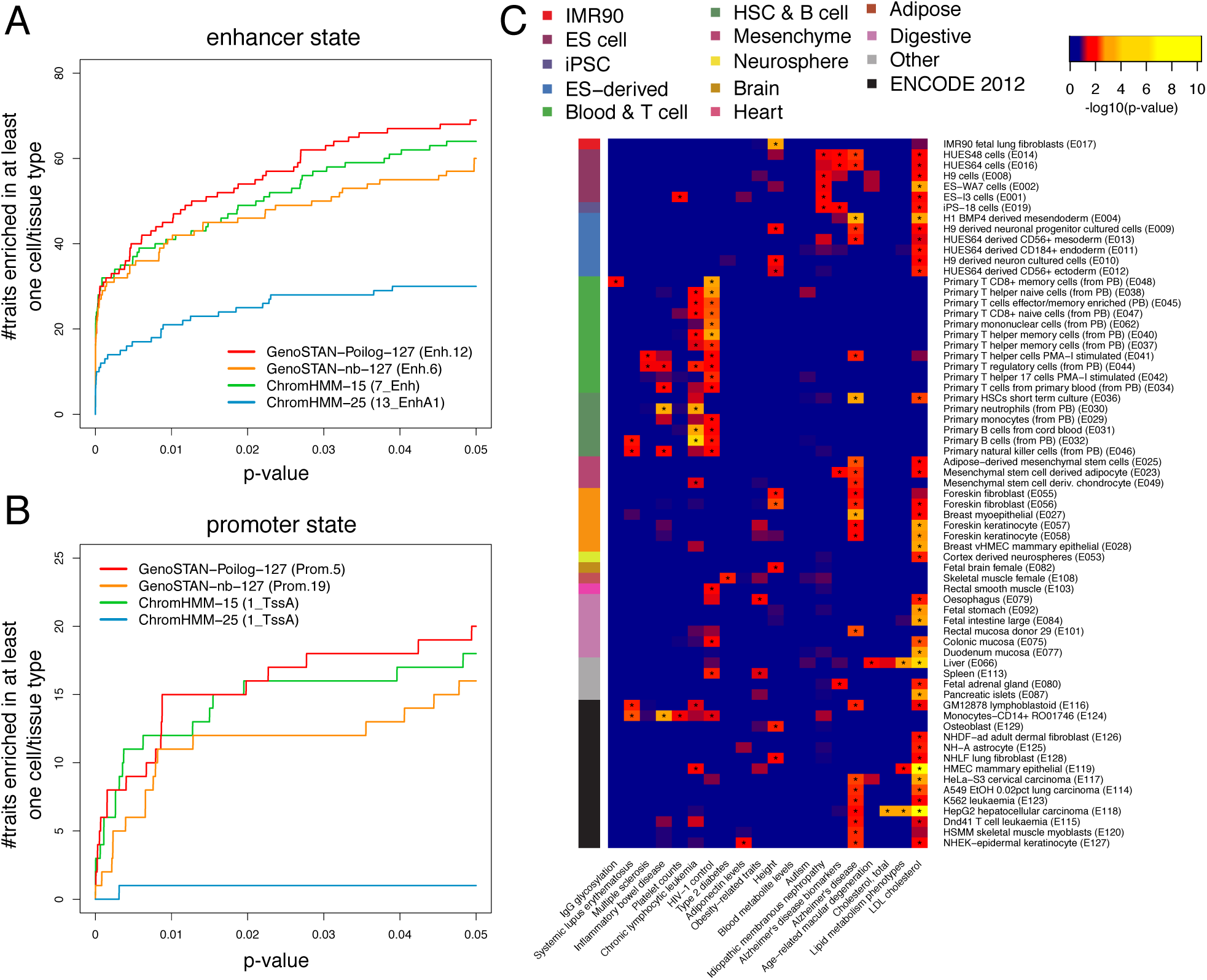
Enrichments of genetic variants associated with diverse traits in enhancers and promoters are specific to the relevant cell types. (A) The number of traits which are enriched in enhancer states in at least one cell type or tissue is plotted for p-values < 0.05. (B) The same as in (A) but for promoters. (C) The heatmap shows the-log10(p-value) of significantly enriched traits in promoter states (GenoSTAN-Poilog-127, p-value < 0.05, marked by ‘*’). P-values were adjusted for multiple testing using the Benjamin-Hochberg correction.

**Supplementary Figure 9:**
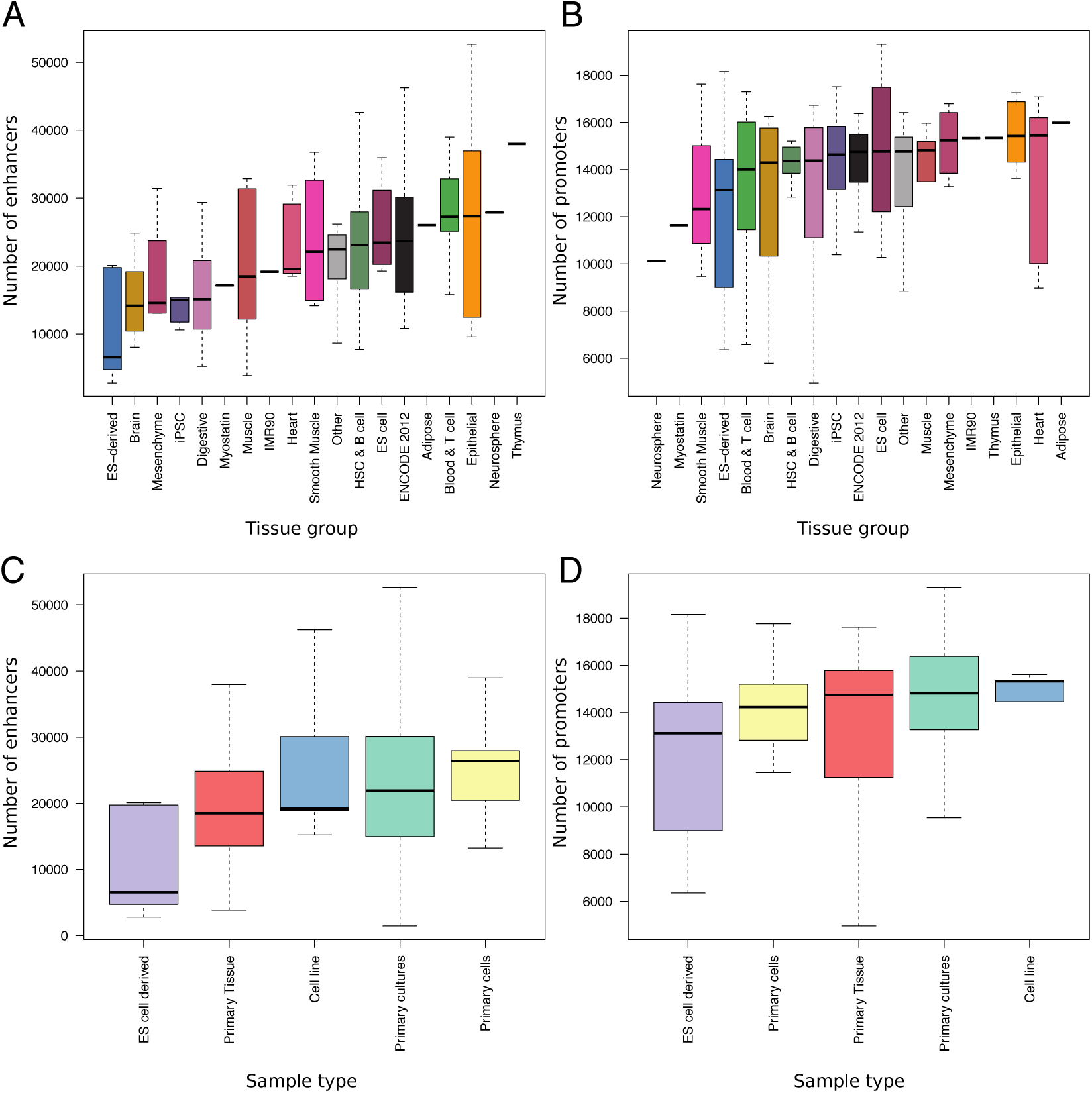
Dependency of number of predicted promoters and enhancers on tissue group and sample type. (A) Number of enhancer states per Roadmap Epigenomics cell/tissue group. (B) The same as in (A) for promoters. (C) Number of enhancer states per Roadmap Epigenomics sample type. (D) The same as in (C) for promoters.

## Supplementary Tables

**Supplementary Table 1:**
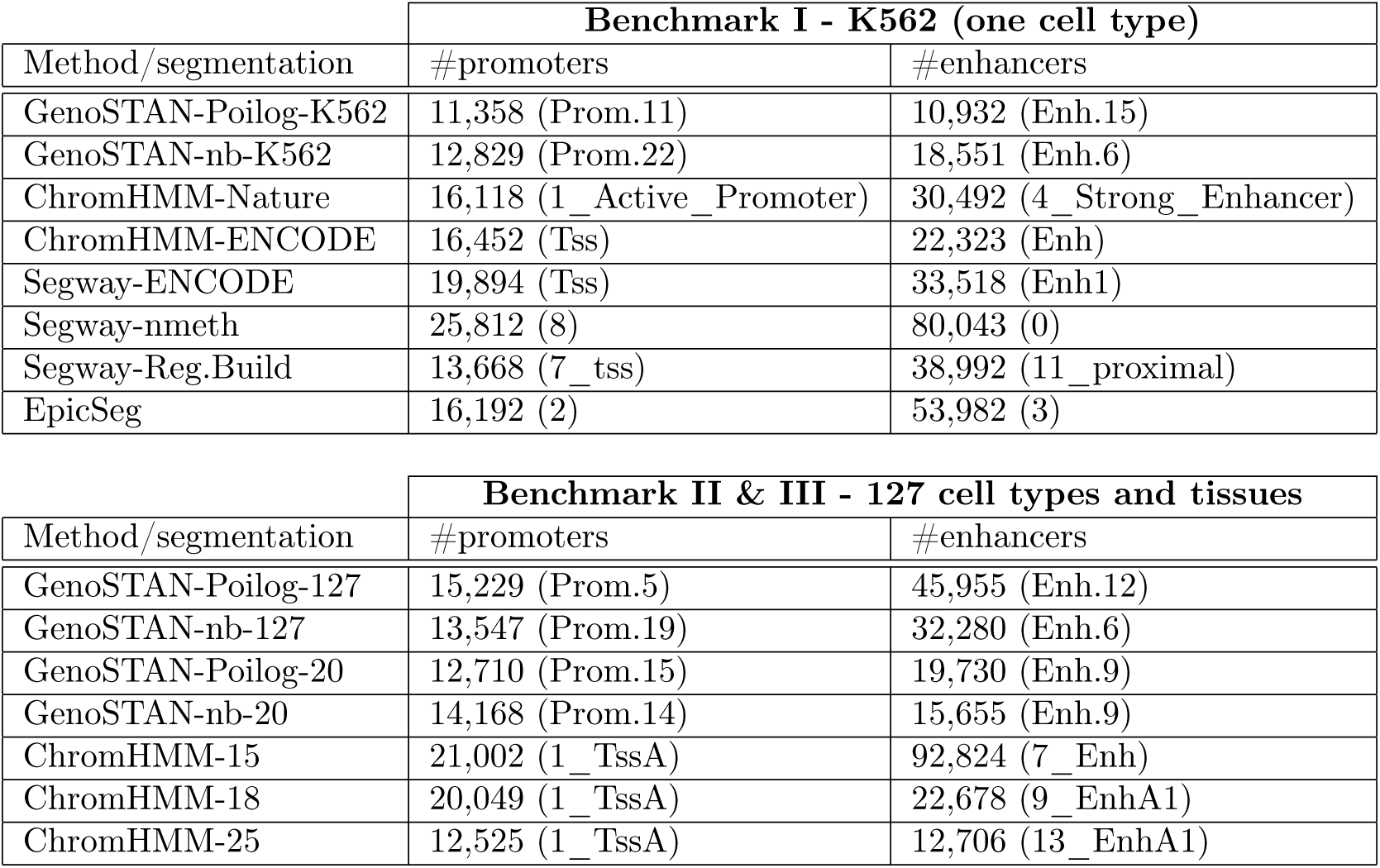
Number of promoter and enhancer states for the chromatin state annotations analyzed in this study. The original state name is given in brackets.

**Supplementary Table 2:** Table showing promoter and enhancers states used to calculate recall of FANTOM5 promoters and enhancers. Two promoter and enhancer states were used for each segmentation, except for the EpicSeg segmentation, which only fitted one enhancer state.

|  | **Benchmark I - K562 (one cell type)** |  |
| --- | --- | --- |
| Method/segmentation | promoter states | enhancer states |
| GenoSTAN-Poilog-K562 | Prom.11, PromW.5 | Enh.15, Enh.2 |
| GenoSTAN-nb-K562 | Prom.16, Prom.22 | Enh.6, Enh.19 |
| ChromHMM-Nature | Tss, TssF | Enh, EnhW |
| ChromHMM-ENCODE | 1_Active_Promoter, 2_Weak_Promoter | 4_Strong_Enhancer, 5_Strong_Enhancer |
| Segway-ENCODE | Tss, PromF | Enh1, Enh2 |
| Segway-nmeth | 8,6 | 0, 13 |
| Segway-Reg.Build | 7_tss, 0_proximal | 1_proximal, 11_proximal |
| EpicSeg | 2 | 3 |

|  | **Benchmark II & III - 127 cell types and tissues** |  |
| --- | --- | --- |
| Method/segmentation | promoter states | enhancer states |
| GenoSTAN-Poilog-127 | Prom.19, Prom.5 | Enh.12, EnhW.9 |
| GenoSTAN-nb-127 | Prom.1, Prom.19 | Enh.6, EnhW.8 |
| GenoSTAN-Poilog-20 | Prom.15, Prom.6 | Enh.9, EnhF.13 |
| GenoSTAN-nb-20 | Prom.14, Prom.21 | Enh.9, EnhF.12 |
| ChromHMM-15 | 1_TssA, 2_PromU | 13_EnhA1, 14_EnhA2 |
| ChromHMM-18 | 1_TssA, 2_TssFlnk | 9_EnhA1, 10_EnhA2 |
| ChromHMM-25 | 1_TssA, 2_TssAFlnk | 7_Enh, 6_EnhG |

## References

[1] Lenhard, B., Sandelin, A. & Carninci, P. Metazoan promoters: emerging characteristics and insights into transcriptional regulation. Nat. Rev. Genet. 13, 233–245 (2012).

[2] Banerji, J., Rusconi, S. & Schaffner, W. Expression of a beta-globin gene is enhanced by remote SV40 DNA sequences. Cell 27, 299–308 (1981).

[3] Kwasnieski, J. C., Fiore, C., Chaudhari, H. G. & Cohen, B. A. High-throughput functional testing of ENCODE segmentation predictions. Genome Res. 24, 1595–1602 (2014).

[4] Kheradpour, P. et al. Systematic dissection of regulatory motifs in 2000 predicted human enhancers using a massively parallel reporter assay. Genome Res. 23, 800–811 (2013).

[5] Arnold, C. D. et al. Genome-wide quantitative enhancer activity maps identified by STARR-seq. Science 339, 1074–1077 (2013).

[6] Shlyueva, D., Stampfel, G. & Stark, A. Transcriptional enhancers: from properties to genome-wide predictions. Nat. Rev. Genet. 15, 272–286 (2014).

[7] Andersson, R. Promoter or enhancer, what’s the difference? Deconstruction of established distinctions and presentation of a unifying model. BioEssays 37, 314–323 (2015).

[8] Ernst, J. & Kellis, M. Discovery and characterization of chromatin states for systematic annotation of the human genome. Nat. Biotechnol. 28, 817–825 (2010).

[9] Hoffman, M. M. et al. Unsupervised pattern discovery in human chromatin structure through genomic segmentation. Nat. Methods 9, 473–476 (2012).

[10] Kleftogiannis, D., Kalnis, P. & Bajic, V. B. Progress and challenges in bioinformatics approaches for enhancer identification. Brief. Bioinformatics [Epub ahead of print] (2015).

[11] Dunham, I. et al. An integrated encyclopedia of DNA elements in the human genome. Nature 489, 57–74 (2012).

[12] Yip, K. Y. et al. Classification of human genomic regions based on experimentally determined binding sites of more than 100 transcription-related factors. Genome Biol. 13, R48 (2012).

[13] Kundaje, A. et al. Integrative analysis of 111 reference human epigenomes. Nature 518, 317–330 (2015).

[14] The blueprint project. www.blueprint-epigenome.eu/.

[15] Andersson, R. et al. An atlas of active enhancers across human cell types and tissues. Nature 507, 455–461 (2014).

[16] Forrest, A. R. et al. A promoter-level mammalian expression atlas. Nature 507, 462–470 (2014).

[17] Kleftogiannis, D., Kalnis, P. & Bajic, V. B. DEEP: a general computational framework for predicting enhancers. Nucleic Acids Res. 43, e6 (2015).

[18] Lee, D., Karchin, R. & Beer, M. A. Discriminative prediction of mammalian enhancers from DNA sequence. Genome Res. 21, 2167–2180 (2011).

[19] Rajagopal, N. et al. RFECS: a random-forest based algorithm for enhancer identification from chromatin state. PLoS Comput. Biol. 9, e1002968 (2013).

[20] Won, K. J. et al. Comparative annotation of functional regions in the human genome using epigenomic data. Nucleic Acids Res. 41, 4423–4432 (2013).

[21] Ernst, J. et al. Mapping and analysis of chromatin state dynamics in nine human cell types. Nature 473, 43–49 (2011).

[22] Hoffman, M. M. et al. Integrative annotation of chromatin elements from ENCODE data. Nucleic Acids Res. 41, 827–841 (2013).

[23] Zerbino, D. R., Wilder, S. P., Johnson, N., Juettemann, T. & Flicek, P. R. The ensembl regulatory build. Genome Biol. 16, 56 (2015).

[24] Kharchenko, P. V. et al. Comprehensive analysis of the chromatin landscape in Drosophila melanogaster. Nature 471, 480–485 (2011).

[25] Rabiner, L. R. A tutorial on hidden Markov models and selected applications in speech recognition. Proc. IEEE 77, 257–286 (1989).

[26] Ernst, J. & Kellis, M. ChromHMM: automating chromatin-state discovery and characterization. Nat. Methods 9, 215–216 (2012).

[27] Heintzman, N. D. et al. Distinct and predictive chromatin signatures of transcriptional promoters and enhancers in the human genome. Nat. Genet. 39, 311–318 (2007).

[28] Mammana, A. & Chung, H. R. Chromatin segmentation based on a probabilistic model for read counts explains a large portion of the epigenome. Genome Biol. 16, 151 (2015).

[29] Zacher, B., Lidschreiber, M., Cramer, P., Gagneur, J. & Tresch, A. Annotation of genomics data using bidirectional hidden Markov models unveils variations in Pol II transcription cycle. Mol. Syst. Biol. 10, 768 (2014).

[30] Heinz, S., Romanoski, C. E., Benner, C. & Glass, C. K. The selection and function of cell type-specific enhancers. Nat. Rev. Mol. Cell Biol. 16, 144–154 (2015).

[31] May, D. et al. Large-scale discovery of enhancers from human heart tissue. Nat. Genet. 44, 89–93 (2012).

[32] Visel, A. et al. ChIP-seq accurately predicts tissue-specific activity of enhancers. Nature 457, 854–858 (2009).

[33] Harrow, J. et al. GENCODE: the reference human genome annotation for The ENCODE Project. Genome Res. 22, 1760–1774 (2012).

[34] Core, L. J. et al. Analysis of nascent RNA identifies a unified architecture of initiation regions at mammalian promoters and enhancers. Nat. Genet. 46, 1311–1320 (2014).

[35] Kim, T.-K. et al. Widespread transcription at neuronal activity-regulated enhancers. Nature 465, 182–187 (2010).

[36] Kvon, E. Z., Stampfel, G., Yanez-Cuna, J.O., Dickson, B. J. & Stark, A. HOT regions function as patterned developmental enhancers and have a distinct cis-regulatory signature. Genes Dev. 26, 908–913 (2012).

[37] Li, H. et al. Functional annotation of HOT regions in the human genome: implications for human disease and cancer. Sci Rep 5, 11633 (2015).

[38] Ernst, J. & Kellis, M. Large-scale imputation of epigenomic datasets for systematic annotation of diverse human tissues. Nat. Biotechnol. 33, 364–376 (2015).

[39] Weinhold, N., Jacobsen, A., Schultz, N., Sander, C. & Lee, W. Genome-wide analysis of noncoding regulatory mutations in cancer. Nat. Genet. 46, 1160–1165 (2014).

[40] Ward, L. D. & Kellis, M. HaploReg v4: systematic mining of putative causal variants, cell types, regulators and target genes for human complex traits and disease. Nucleic Acids Res. 44, D877–881 (2016).

[41] Trynka, G. et al. Chromatin marks identify critical cell types for fine mapping complex trait variants. Nat. Genet. 45, 124–130 (2013).

[42] Lindblad-Toh, K. et al. A high-resolution map of human evolutionary constraint using 29 mammals. Nature 478, 476–482 (2011).

[43] Welter, D. et al. The NHGRI GWAS Catalog, a curated resource of SNP-trait associations. Nucleic Acids Res. 42, D1001–1006 (2014).

[44] Ramskold, D., Wang, E. T., Burge, C. B. & Sandberg, R. An abundance of ubiquitously expressed genes revealed by tissue transcriptome sequence data. PLoS Comput. Biol. 5, e1000598 (2009).

[45] Ernst, J. & Kellis, M. Interplay between chromatin state, regulator binding, and regulatory motifs in six human cell types. Genome Res. 23, 1142–1154 (2013).

[46] Gerstein, M. B., Kundaje, A., Hariharan, M., Weissman, S. M. & Snyder, M. Architecture of the human regulatory network derived from ENCODE data. Nature 489, 91–100 (2012).

[47] Wang, J. et al. Sequence features and chromatin structure around the genomic regions bound by 119 human transcription factors. Genome Res. 22, 1798–1812 (2012).

[48] Org, T. et al. Scl binds to primed enhancers in mesoderm to regulate hematopoietic and cardiac fate divergence. EMBO J. 34, 759–777 (2015).

[49] Ihaka, R. & Gentleman, R. R: A Language for Data Analysis and Graphics. J. Comp. Graph. Stat. 5, 299–314 (1996).

[50] Gentleman, R. C. et al. Bioconductor: open software development for computational biology and bioinformat-ics. Genome Biol. 5, R80 (2004).

[51] Zentner, G. E., Tesar, P. J. & Scacheri, P. C. Epigenetic signatures distinguish multiple classes of enhancers with distinct cellular functions. Genome Res. 21, 1273–1283 (2011).

[52] Rabani, M. et al. High-resolution sequencing and modeling identifies distinct dynamic RNA regulatory strategies. Cell 159, 1698–1710 (2014).

[53] Miller, C. et al. Dynamic transcriptome analysis measures rates of mRNA synthesis and decay in yeast. Mol. Syst. Biol. 7, 458 (2011).

[54] Churchman, L. S. & Weissman, J. S. Nascent transcript sequencing visualizes transcription at nucleotide resolution. Nature 469, 368–373 (2011).

[55] Rajagopal, N. et al. High-throughput mapping of regulatory DNA. Nat. Biotechnol. 34, 167–174 (2016).

[56] Cook, J. D. Notes on the negative binomial distribution www.johndcook.com/negative_binomial.pdf.

[57] Bulmer, M. G. On Fitting the Poisson Lognormal Distribution to Species-Abundance Data. Biometrics 30, 101–110 (2011).

[58] Grotan, V. & Engen, S. poilog: Poisson lognormal and bivariate Poisson lognormal distribution (2008). R package version 0.4.

[59] Dillies, M. A. et al. A comprehensive evaluation of normalization methods for Illumina high-throughput RNA sequencing data analysis. Brief. Bioinformatics 14, 671–683 (2013).

[60] Langmead, B. & Salzberg, S. L. Fast gapped-read alignment with Bowtie 2. Nat. Methods 9, 357–359 (2012).

[61] Li, H. et al. The Sequence Alignment/Map format and SAMtools. Bioinformatics 25, 2078–2079 (2009).

[62] Birney, E. et al. Identification and analysis of functional elements in 1% of the human genome by the ENCODE pilot project. Nature 447, 799–816 (2007).

[63] Lawrence, M., Gentleman, R. & Carey, V. rtracklayer: an r package for interfacing with genome browsers. Bioinformatics 25, 1841–1842 (2009).

[64] Lawrence, M. et al. Software for computing and annotating genomic ranges. PLoS Comput. Biol. 9, e1003118 (2013).

